# Tau conformers in FTLD-MAPT undergo liquid-liquid phase separation and perturb the nuclear envelope

**DOI:** 10.1101/2020.07.04.187997

**Authors:** Sang-Gyun Kang, Zhuang Zhuang Han, Nathalie Daude, Emily McNamara, Serene Wohlgemuth, Jiri G. Safar, Sue-Ann Mok, David Westaway

## Abstract

Recent studies show that a single *MAPT* gene mutation can promote alternative tau misfolding pathways engendering divergent forms of frontotemporal dementia and that under conditions of molecular crowding, the repertoire of tau forms can include liquid-liquid phase separation (LLPS). We show here that following pathogenic seeding, tau condenses on the nuclear envelope (NE) and disrupts nuclear-cytoplasmic transport (NCT). Interestingly, NE fluorescent tau signals and small fluorescent inclusions behaved as demixed liquid droplets in living cells. Thioflavin S-positive intracellular aggregates were prevalent in tau-derived inclusions with a size bigger than 3 μm^2^, indicating that a threshold of critical mass in the liquid state condensation may drive liquid-solid phase transitions. Our findings indicate that tau undergoing LLPS is more toxic amongst a spectrum of alternative conformers; LLPS droplets on the NE that disrupt NCT serve to trigger cell death and can act as nurseries for fibrillar structures abundantly detected in end-stage disease.

## Introduction

Intracellular inclusions of microtubule-associated protein tau are the pathological hallmark of tauopathies including frontotemporal lobar degenerations (FTLDs) and Alzheimer’s disease (AD) (Gotz, Halliday, & Nisbet, 2019; T. Guo, Noble, & Hanger, 2017). Tau is encoded by the *MAPT* gene and expressed mainly in neurons as six different isoforms, depending on neuronal types and maturation stages (T. Guo et al., 2017; Wang & Mandelkow, 2016). Tau stabilizes and maintains the architecture of microtubules and axonal integrity of neurons, in which tau is in a dynamic equilibrium between a microtubule-bound and cytoplasmic free state (T. Guo et al., 2017; X. Zhang et al., 2017). The conformational change of monomeric soluble tau into other conformers that include hyperphosphorylated oligomers, paired helical filaments (PHFs) and fibrillized tau is thought to contribute to neuronal toxicity and cell death (Gotz et al., 2019; T. Guo et al., 2017; Wang & Mandelkow, 2016). We recently reported that even the same germline mutation, *MAPT*-P301L, generates distinct tau conformers as appraised by conformation-dependent immunoassay (CDIs) and conformational stability assays (CSAs) (Daude et al., 2020). The diverse and evolving repertoire of tau conformers that includes four CSA profiles in mice (CSA Types 1-4) was postulated as the origin of neuropathological and biochemical heterogeneity of FTLD with tau immunoreactive inclusions (FTLD-tau) (Daude et al., 2020; Gotz et al., 2019). Moreover, in frontotemporal dementia (FTD), a neurological diagnosis that is associated with the neuropathological diagnosis of FTLD, CSA Types were correlated with clinical disease variants. However, this being said, the cellular events that draw a line from protein conformation to neurological dysfunction are not well understood.

Nuclear localization of tau has been observed and suggested to facilitate genome surveillance under conditions of cellular stresses (Bukar Maina, Al-Hilaly, & Serpell, 2016). In neurodegenerative disorders including FTD, Huntington’s disease, Parkinson disease and amyotrophic lateral sclerosis (ALS), disruption of nuclear-cytoplasmic transport (NCT) has been proposed as a toxic mechanism mediated by abnormally aggregated proteins (Grima et al., 2017; Jiang et al., 2016; Jovicic et al., 2015; Woerner et al., 2016; K. Zhang et al., 2018; K. Zhang et al., 2015). Nuclear pore complexes (NPCs), which are one of the largest macromolecular assemblies found in eukaryotic cells, reside in the nuclear envelope (NE) and mediate NCT of various nuclear proteins and RNAs (Clarke & Zhang, 2008; Guttinger, Laurell, & Kutay, 2009; Timney et al., 2016). These cellular components, as well as lamin proteins that contribute to the lamina of the NE, may have an intrinsic jeopardy to accumulating damage in chronological aging as they have remarkably low rates of turnover (Toyama et al., 2013). For tau, it is accepted that alterations in the physiological properties resulting from post-translational modifications, conformational changes and/or pathogenic mutations, can lead to mis-localization and formation of inclusions in neuronal cell bodies (Gotz et al., 2019; T. Guo et al., 2017). More recently, there has been a focus upon whether tau inclusions cause an impairment of NCT, following from sequestration of nucleoporins (NUPs) (Eftekharzadeh et al., 2018) and nuclear deformation (Paonessa et al., 2019) (both *in vitro* and *in vivo*), incurring toxic consequences. It is possible that these newly documented changes in tau and nuclear proteins may intersect with discoveries in a third axis of work.

Membraneless organelles (MLOs) formed by a phase separation process (see below) have been highlighted as active bioreactors regulating cell signaling, protein synthesis and various biological reactions against environmental stresses (Alberti, Gladfelter, & Mittag, 2019; Brangwynne, 2013; Ryan & Fawzi, 2019; Shin & Brangwynne, 2017). Rapid and reversible phase transition of MLOs simultaneously presents a fascinating chemical change and opens up a new frontier in pathogenesis, relating to the cellular and molecular impacts of these assemblies (Brangwynne, 2013; Ryan & Fawzi, 2019). Multivalent polymers, especially proteins containing low complexity domains (LCDs) and RNA molecules, bind to each other and condense as liquid droplets, a process termed liquid-liquid phase separation (LLPS) that has been thought to regulate MLOs (Alberti et al., 2019; Ryan & Fawzi, 2019; Shin & Brangwynne, 2017). MLOs need to be assembled as functional condensed droplets and be disassembled by quality control processes within a confined biological time-scale if irreversible conformational changes are to be avoided (Nedelsky & Taylor, 2019; Patel et al., 2015). In ALS/FTD, loss-of-function mutations in LCDs and/or RNA recognition motifs (RRMs) are responsible for neurotoxicity by disrupting the dynamics of MLOs (Ryan & Fawzi, 2019). Pathogenic mutations in DNA-binding protein 43 (TDP43), heterogeneous nuclear ribonucleoprotein A1 (hnRNPA1) and fused in sarcoma (FUS) altered biophysical properties of MLOs from reversible metastable liquid condensates to irreversible persistent fibrous aggregates (Kim et al., 2013; Mann et al., 2019; Murakami et al., 2015; Patel et al., 2015). Toxic dipeptide repeat (DPRs) proteins are produced from a hexanucleotide repeat expansion in C9ORF72, which is the most common cause of ALS/FTD (Jovicic et al., 2015; K. Zhang et al., 2015) and the interactions of DPRs with proteins harboring LCDs or RRMs disturb multiple MLOs such as nucleoli, NPCs and stress granules (SGs) (Lee et al., 2016). For tau, recent studies have shown that this intrinsically disordered protein, although lacking predicted LCDs and RRMs, nonetheless has some propensity to undergo LLPS (Ambadipudi, Biernat, Riedel, Mandelkow, & Zweckstetter, 2017; Boyko, Qi, Chen, Surewicz, & Surewicz, 2019; Singh, Xu, Boyko, Surewicz, & Surewicz, 2020; Vega, Umstead, & Kanaan, 2019; Wegmann et al., 2018; X. Zhang et al., 2017).

In studies here we now demonstrate that liquid phase condensation of tau occurs in living cells and that this effect derives from gain-of-function properties of FTLD-MAPT mutations in the repeat domain (i.e., P301L or P301L+V337M). In contrast to the loss-of-function mutations in TDP43, hnRNPA1, FUS and C9ORF72 (DPRs) that downgrade a physiological, protective form of LLPS (Kim et al., 2013; Lee et al., 2016; Mann et al., 2019; Murakami et al., 2015; Patel et al., 2015), disease-causing mutations in tau facilitate LLPS assemblies that sequester NUPs from NPCs and hence are toxic by virtue of impeding vital NCT. Because of these toxic cellular effects, LLPS tau can be seen as an important entity within a spectrum of tau conformers defined by chemical denaturation (Daude et al., 2020).

## Results

### Nuclear architecture in FTLD-tau

We have previously reported a slow model of a primary tauopathy, FTLD-MAPT; aged mice from the TgTau^P301L^ line can show heterogeneity in histopathological presentations and types of trypsin-resistant cores of tau, phenomena which are likely related to the variations in clinical phenotypes seen in FTLD-MAPT-P301L patients (Borrego-Ecija et al., 2017; Daude et al., 2020; Eskandari-Sedighi et al., 2017; Murakami et al., 2006). To investigate tau-associated nuclear distortion, nuclear lamina in post-mortem cerebral cortex of both FTLD-MAPT-P301L patients and TgTau^P301L^ mice were probed by lamin B1 immunostaining. Our resources included brain tissue from ten Iberian FTLD-MAPT-P301L patients (Borrego-Ecija et al., 2017), that likely derive from a common ancestor (Palencia-Madrid et al., 2019). These P301L cases, characterized previously for tau pathology and a confirmed absence of confounding proteinopathies (Borrego-Ecija et al., 2017; Daude et al., 2020), were augmented by a number of controls including AD cases, FTD with progranulin mutations, ALS cases and non-demented controls. Although analyses by others have remarked upon nuclear clefts as a feature of FTLD-MAPT (Paonessa et al., 2019), when examining the nuclei of dentate gyrus (DG) neurons this finding also applied to other clinical entities, being abundant within three ALS cases, two progranulin mutation carriers and in one non-demented control (**Table 1** and **Figure 1**). While there was a trend for lower ages in the P301L group, this did not reach significance and this alteration was thus considered to be age-related and not disease-related. Thus, along with other analyses (Molina-Porcel et al., 2019), the hypothesis for a relationship between nuclear clefts and the specific pathogenic processes of FTLD-MAPT was not supported, prompting consideration of other nuclear alterations caused by the presence of misfolded tau isoforms. Using anti-lamin B1 antibodies to stain the nuclear lamina, we assessed potential distinctions between FTLD-MAPT-P301L cases versus control samples (**Figure 1a** to **1d**). Discounting occasional nuclear clefts (**Table 1, Figure 1b** and **1e**) also present in other diseases, several distinctions were noted, which included: variations in staining intensity on the margins of normally shaped nuclei (**Figure 1d**), nuclei with angled margins and non-uniform lamin B staining (**Figure 1f**) and cells with granular and apparently spherical immunostained structures in the cytoplasm (**Figure 1g** and **1h**). Considering the mouse FTLD-MAPT-P301L model, nuclear clefts were present in both aged TgTau^P301L^ and non-Tg mice (**Figure 1i** to **1l**), but we observed nuclei with angled margins and non-uniform lamin B1 staining in the cortex and DG for Tg mice as well as incomplete staining of nuclear lamina (**Figure 1m** to **1p**). Juxtanuclear tau signals and punctate cytoplasmic signals were present in both FTLD-MAPT-P301L cases and TgTau^P301L^ mice, along with some nuclear margins decorated by interspersed circular areas of tau staining (**Figure 1q)**. Indeed, double-staining experiments yielded an apparently reciprocal pattern of staining where tau signals along the nuclear margins were matched by dimmed areas of lamin B staining (**Figure 1q**). These data suggested an exchange or swapping phenomenon affecting proteins on or adjacent to the nuclear envelope.

**Figure 1.**
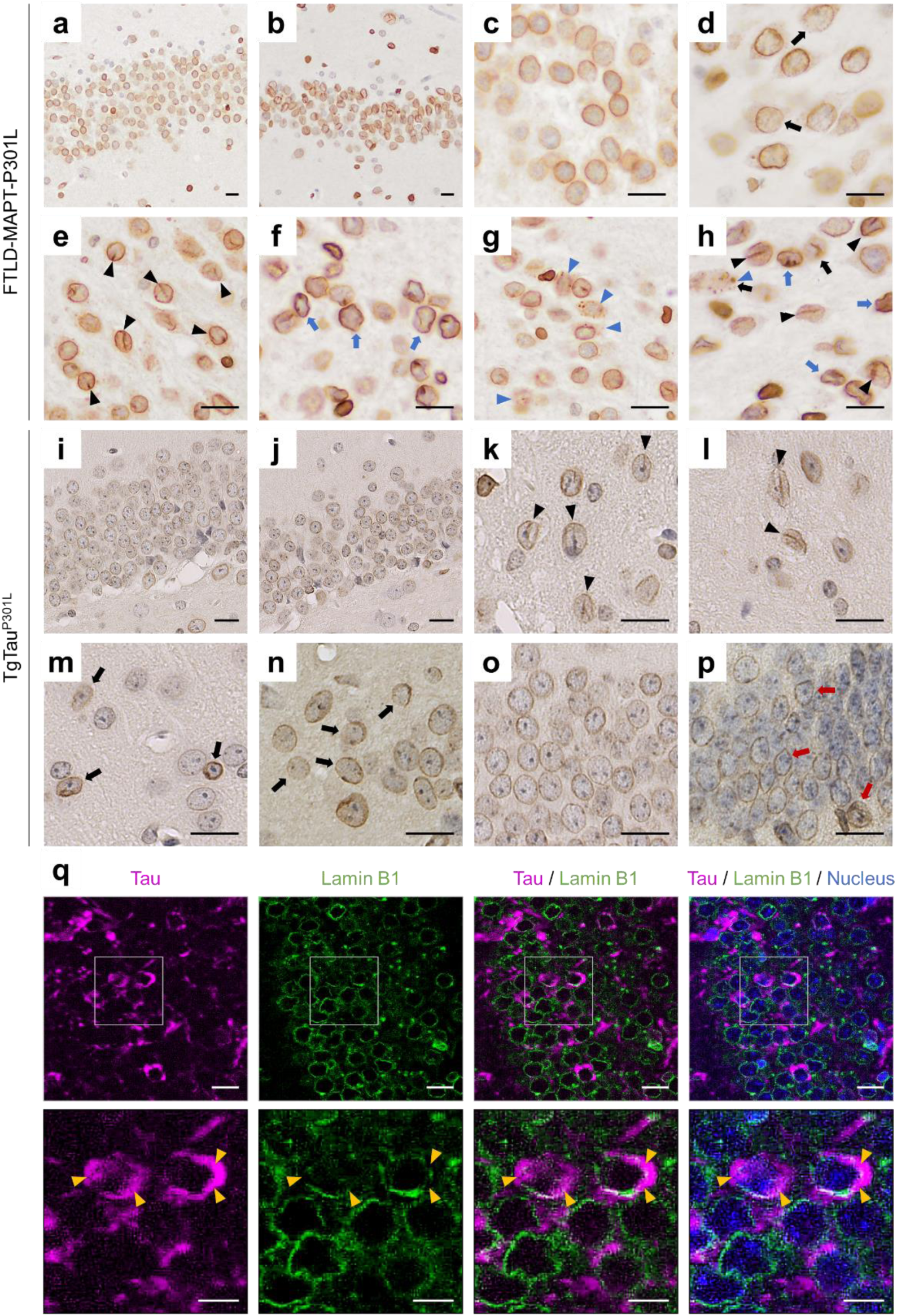
Deformation of neuronal nuclear envelopes in FTLD-tau. Lamin B1 staining for nuclear lamina delineated the nuclear membrane of granular neurons in the dentate gyrus (DG) of the hippocampus in a control (**a**). In contrast, neuronal loss and nuclear morphological changes were evident in the DG of a MAPT-P301L mutation carrier (**b**). In comparison with the control (**c**), lamin B1 staining was thinner and incomplete in the MAPT-P301L patient (**d**, black arrows). Nuclear deformations including intranuclear clefts (**e**, black arrowheads) and angular nuclear morphologies (**f**, blue arrows) were observed in the patient with MAPT-P301L mutation. Some neurons appeared with cytoplasmic granular staining of lamin B1 (**g**, blue arrowheads) in the region where incomplete lamin B1 staining (black arrow), intranuclear clefts (black arrowheads) and angular nuclei (blue arrows) were found (**h**). Compared to the non-Tg mice (**i**), TgTau^P301L^ showed loss of granular neurons in the DG (**j**). Intranuclear clefts (black arrowheads) were observed in the thalamus of aged non-Tg (**k**) and in TgTau^P301L^ mice (**l**). Variations in staining intensities of lamin B1 (black arrows) in the frontal cortex (**m**, thicker) and the DG (**n**, thinner and incomplete) of TgTau^P301L^. Unlike the control (**o**), angular nuclear morphologies were found in the DG of the TgTau^P301L^ (**p**). **q**. Immunofluorescent staining of phosphorylated tau (magenta, AT8) and lamin B1 (green) revealed that areas of discontinued nuclear membrane were overlapped with tau deposits (yellow arrowheads) in TgTau^P301L^. Nuclei were counterstained with DAPI (blue). Scale bars, 20 μm and 10 μm in the boxed images.

**Table 1.** Nuclear deformation of granular neurons in the dentate gyrus of human patients with FTLD-MAPT or other neurodegenerative disorders

| Clinical diagnosis | Age at death | Sex | Nuclear morphology based on lamin B1 staining |  |  |  | Phospho-tau |
| --- | --- | --- | --- | --- | --- | --- | --- |
|  |  |  | Cleft | Cytopl | Discont | Angular |  |
| Control | 31 | M | 0.5 | 0 | 0 | 0.5 | 0 |
| Control | 70 | M | 1 | 0 | 0 vs UK | 2 | 0 |
| Control | 76 | M | 2 | 0 | 0 | 1 | 1+ |
| Control | 78 | M | 3 | 0 | 0 | 2 | 1+ |
| Control | 81 | F | 1 | 0 | 0 vs UK | 1 | 0 |
| AD | 72 | F | 0.5 | 0 | 0 | 0.5 | 1+ |
| AD | 92 | F | 3 | 0.5 | 0.5 | 3 | 2+ |
| *GRN* | 60 | M | 3 | 0.5 | 0.5 | 2 | 0 |
| *GRN* | 64 | M | 3 | 0 | 0.5 | 3 | 1+ |
| ALS | 50 | M | 3 | 0 | 0 | 2 | 0 |
| ALS | 50 | M | 3 | 0 | 0 | 1 | 0 |
| ALS | 51 | M | 2 | 0 | 0 | 1 | 0 |
| ALS | 63 | M | 3 | 0 | 0.5 | 1 | 0 |
| ALS | 64 | M | 2 | 0 | 0 | 1 | 0 |
| ALS/FTD | 59 | M | 2 | 0.5 | 0.5 | 1 | 0 |
| FTLD-P301L | 49 | M | 3 | 2 | UK | 2 | ND |
| FTLD-P301L | 52 | M | 2 | 2 | 1 | 3 | 3+ |
| FTLD-P301L | 53 | M | 3 | 3 | 1 | 3 | 3+ |
| FTLD-P301L | 56 | M | 3 | 3 | 2 | 2 | 3+ |
| FTLD-P301L | 58 | M | 2 | 2 | 1 | 1 | 3+ |
| FTLD-P301L | 58 | M | 3 | 3 | 2 | 2 | 3+ |
| FTLD-P301L | 61 | F | 2 | 1 | 1 | 1 | 3+ |
| FTLD-P301L | 63 | F | 2 | 2 | 1 | 2 | 3+ |
| FTLD-P301L | 72 | M | 3 | 3 | 2 | 3 | 3+ |
| FTLD-P301L | 75 | M | 3 | 1 | 0.5 | 3 | 3+ |

Table after (Borrego-Ecija et al., 2017), arranged by clinical diagnosis: control, non-neurological diseases; AD, Alzheimer disease; *GRN*, mutations in the progranulin gene; ALS, amyotrophic lateral sclerosis; ALS/FTD, ALS and frontotemporal dementia; FTLD-P301L, frontotemporal lobar degeneration with P301L mutation in the *MAPT* gene. M, male; F, female. Cleft, nuclear cleft; Cytopl, cytoplasmic granular lamin B stain; Discont, discontinuous nuclear edge; Angular, angled nuclear envelope. UK, unknown; ND, not detected. Scoring criteria for tau deposits as per (Borrego-Ecija et al., 2017; Daude et al., 2020).

### Tau inclusions accumulate on the nuclear envelope

To further investigate tau aggregation and its putative cytotoxicity, brain homogenates derived from TgTau^P301L^ mice exhibiting pathological signs of neurological disease were seeded into two distinct tau reporter cells (Eskandari-Sedighi et al., 2017; Kaufman et al., 2016; Sanders et al., 2014). Firstly, we used human embryonic kidney 293 cells (HEK293) expressing yellow fluorescent protein (YFP) fused the four-repeat domain (4R) of human tau with aggregation prone mutations (P301L/V337M), 4RD-YFP reporter cells (4RD-YFP P301L/V377M) (Sanders et al., 2014). Secondly, we also used HEK293 cells expressing a doxycycline-inducible green fluorescent protein (GFP) fused full-length human tau (0N4R) with aggregation prone mutation (P301L), GFP-0N4R reporter cells (Dox:GFP-0N4R P301L). The GFP-0N4R form of tau (a 66 kDa species) was observed in cytoplasm as expected, while the 4RD-YFP tau (a 45 kDa species) yielded signals spread throughout the cell body to include the nucleus (**Figure 2a** and **2b**); the latter may be due to passive macromolecular diffusion through NPCs which decreases beyond a 30-60 kDa size threshold (Timney et al., 2016).

**Figure 2.**
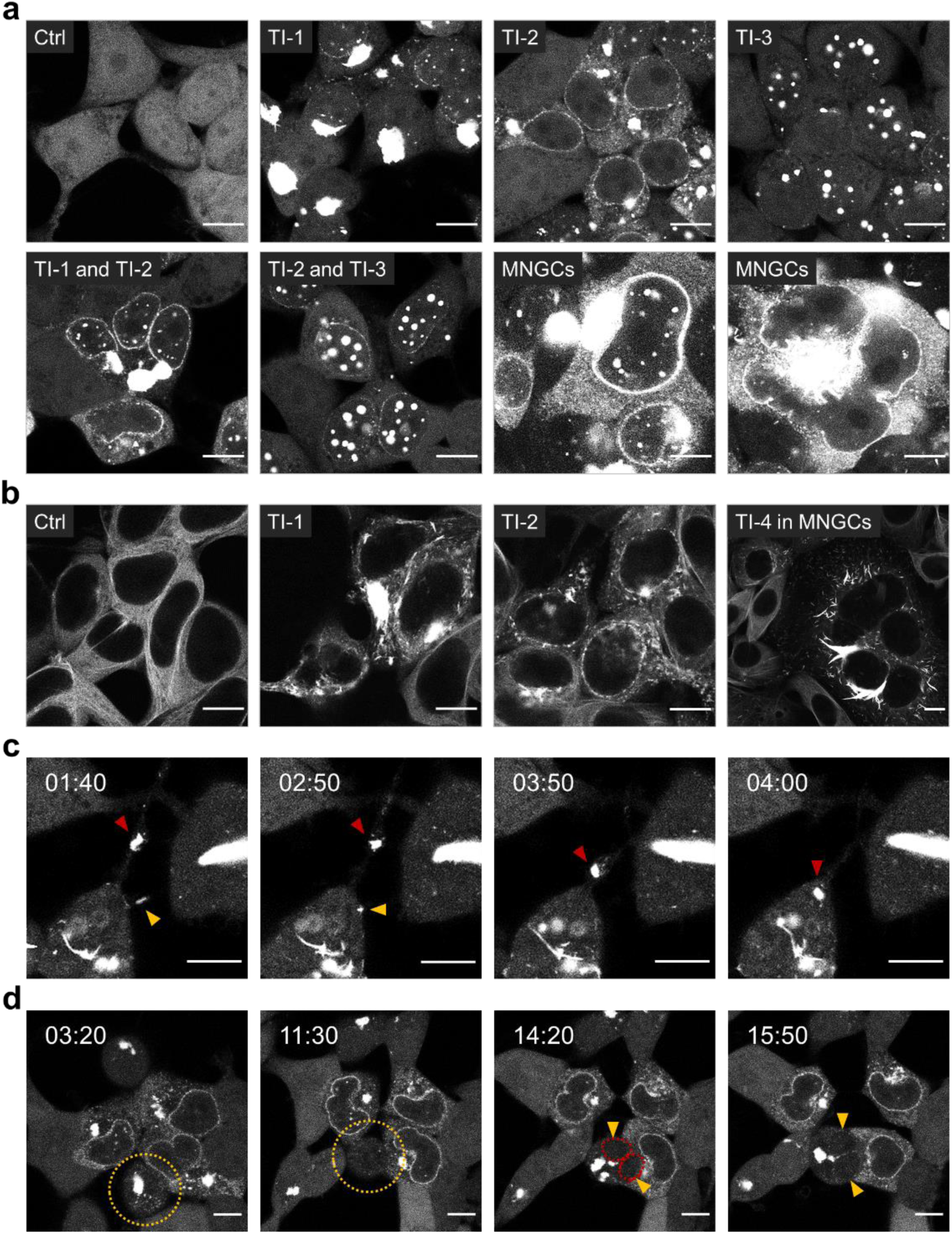
Heterogeneous morphology of tau inclusions in tau reporter cells. **a** and **b.** Tau reporter cells (**a**, 4RD-YFP P301L/V377M; **b**, Dox:GFP-0N4R P301L) were seeded with brain homogenate of aged TgTau^P301L^ including CSA Type 2 tau conformers and imaged at 6 days post seeding. Diverse tau inclusion (TI) morphologies were observed; a large mass of aggregated tau with no specific pattern (amorphous, TI-1), juxtanuclear and nuclear membrane inclusions (nuclear envelope, NE, TI-2), granular inclusions (speckles, TI-3), and threads (TI-4). Some cells showed mixed TI morphologies appearing as TI-1 and TI-2, or TI-2 and TI-3 simultaneously. Multinucleated giant cells (MNGCs) were characterized by apparent NE inclusions with increased cytoplasmic tau signals (**a**) and thread shapes (**b**). **c** and **d**. Live cell imaging analysis of the seeded reporter cells (4RD-YFP P301L/V377M). Time-lapse images were collected at 6 days post seeding by recording photographs for 16 hours at one frame every 10 min (1/10 frame/min). **c**. Cell-to-cell spread of tau inclusions through tunneling nanotube-like protrusion of plasma membrane (both red and yellow arrowheads). **d.** Multinucleated cells emerged through a failure in cell division (yellow arrowheads). Ctrl, control cells seeded with the brain homogenate of non-Tg mice. Scale bar, 10 μm.

Confirming and extending previous analyses (Daude et al., 2020), fluorescent signatures included large tau inclusions in cytoplasm (amorphous, TI-1), discontinuous perimeter signals along with the nuclear edges (nuclear envelope, NE, TI-2), small bead shapes with various sizes most likely seen in nucleus (speckles, TI-3) and cytoplasmic fibril-like strip forms (threads, TI-4) (**Figure 2a** and **2b**, **Supplementary Figure 1**). Seeded tau reporter cells occasionally appeared with a complex of mixed morphologies (TI-1 and TI-2, or TI-2 and TI-3), or as multinucleated giant cells (MNGCs), characterized by bright NE with increased cytoplasmic tau signals or various threads shaped inclusions (**Figure 2a** and **2b**). Following these baseline descriptions of fixed cells, live cell imaging was then undertaken to investigate dynamic aspects of tau inclusion formation. These analyses revealed cell-to-cell spread and, also, mitosis of fluorescence-positive seeded cells often resulted in both daughter cells being positive for tau inclusions (**Supplementary Figure 2a** and **Supplementary Movie 1**). We noted that tau inclusions within cell debris were adsorbed by adjacent cells and fused with others, which produced larger inclusions (**Figure 2c** and **Supplementary Movie 2**, **Supplementary Figure 2b** and **Supplementary Movie 3**) as being consistent with the previous report that dynamic structure of tau aggregates undergo “fusion” and “fission” in stable cell lines expressing full-length human tau T40 (2N4R) carrying the P301L mutation with a GFP tag (T40/P301L-GFP) (J. L. Guo et al., 2016). Moreover, live cell imaging analyses indicated that MNGCs resulted from a failure in cell division (**Figure 2d** and **Supplementary Movie 4**); as reported by others (Caneus et al., 2018), mitotic abnormalities, chromosome mis-segregation, and aneuploidy were observed in transgenic mice expressing the human P301S FTLD-MAPT mutation.

Among the four fluorescent morphologies observed in transduced cells, NE tau inclusions (TI-2) were prominent when seeding reporter cells with brain extracts assigned with a CSA profile called Type 2; this conformational profile for aggregated tau was found in TgTau^P301L^ mice or in frontal cortex extracts from FTLD-MAPT-P301L patients presenting as a behavioral variant of FTD with memory impairment (bvFTD*) (Daude et al., 2020). To allow more detailed biochemical and cell biological investigations of these NE tau inclusions, we seeded 4RD-YFP reporter cells (P301L/V377M) with a CSA Type 2 brain homogenate (Daude et al., 2020) and established a single cell clone by limiting dilution, designated ES1. Interestingly, these ES1 clonal cells exhibited all the aforementioned tau inclusion morphologies described in **Figure 2a**, as well as occasional mixed morphological phenotypes and multinucleated cells (**Figure 3a**). To exclude the occurrence of non-clonal cell isolates surviving the limiting dilution procedure, we re-cloned the ES1 cells by another round of limiting dilutions. Six new single cell clones were obtained, but these were still not obviously distinguishable from the parental ES1 cells. Thus, the single cell clones exhibited the same heterogeneous inclusion phenotypes (**Supplementary Figure 3a**) and the same size of protease-resistant core following limited proteolytic digestion (**Supplementary Figure 3b** to **3e**), suggesting this grouping of phenotypic properties reflect an intrinsic property or capability of the misfolded tau species within this cell clone.

**Figure 3.**
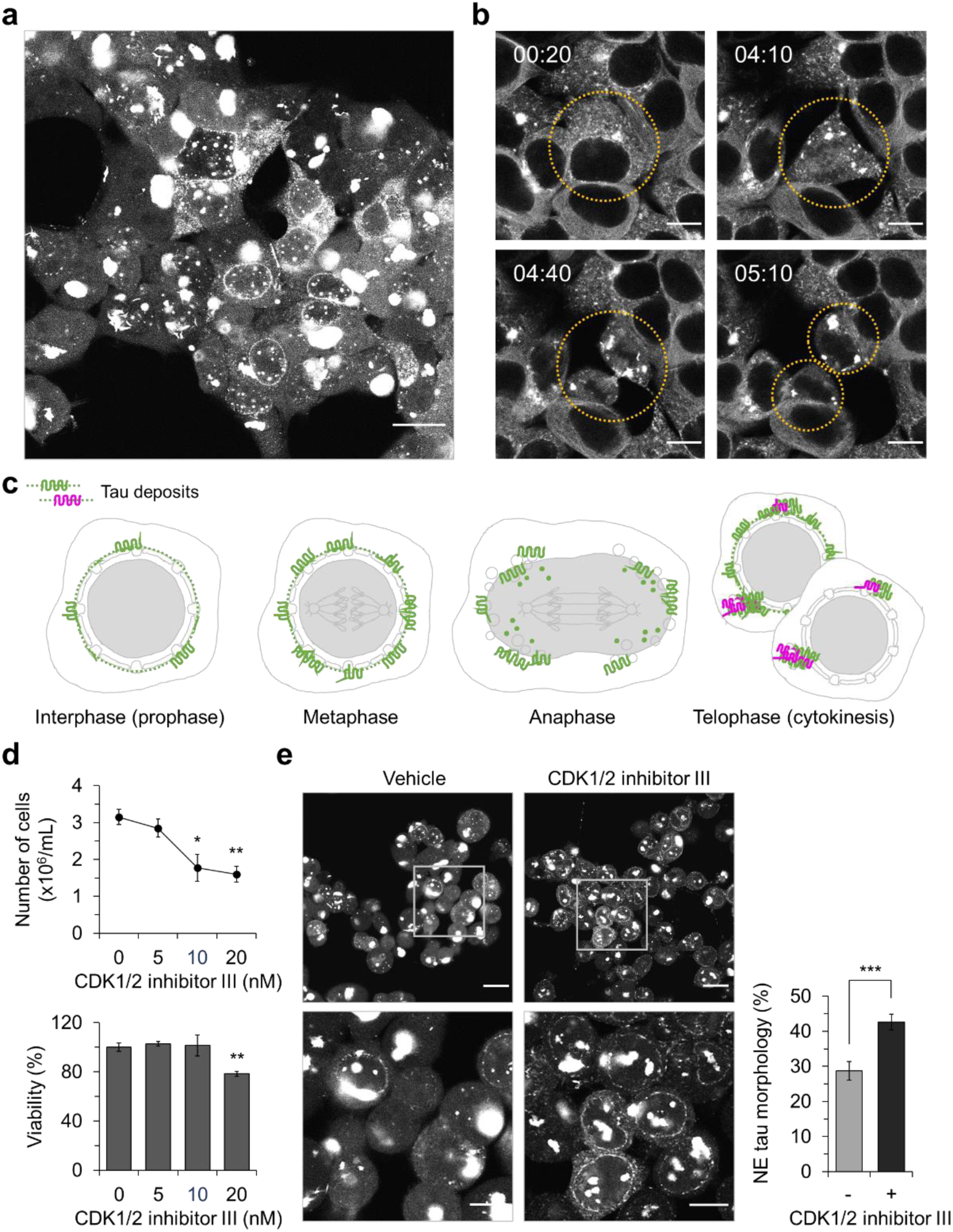
Tau inclusions in the clonal subline on the nuclear envelope. **a**. Tau reporter cells (4RD-YFP P301L/V377M) were seeded with brain homogenates of CSA Type 2 TgTau^P301L^ and subcloned (ES1). The ES1 exhibited heterogenous morphology of tau inclusions. Scale bar, 20 μm. **b**. Live cell imaging as per Figure 2 revealed that tau inclusions in the seeded reporter cells (Dox:GFP-0N4R P301L) underwent morphological changes. Cells were imaged every 10 min for 16 hours. Scale bar, 10 μm. **c**. A schematic of morphological changes of NE tau inclusions. Upon tau seeding, cellular tau starts to condense and recruit on the NE during interphase to prophase. The NE tau loses their morphology since disassembly of the NE commences at metaphase and NE components are dispersed throughout the cytoplasm at anaphase. During telophase, reassembly of the NE is completed, while tau combine with each other and appear as heterogeneous morphologies. **d**. Cell cycle arrest in ES1 cells. CDK1/2 inhibitor III blocked proliferation of ES1 cells with no reduction in viability (n=4). **e**. CDK1/2 inhibitor III increased the number of cells showing NE tau inclusions compared to the vehicle (DMSO) treatments. Numbers of cells with NE tau morphology were counted from 12 and 9 different areas of the cover slip for CDK1/2 inhibitor III (total 781 cells) and vehicle treatments (total 600 cells), respectively. Scale bar, 20 μm and 10 μm in the boxed images. Error bars represent SEM. *p < 0.05 and **p < 0.01 in comparison with the controls.

### Cell cycle effects, nuclear inclusions and cytotoxicity

Quite remarkably, closer analysis of non-multinuclear ES1 cells by live cell imaging analysis revealed dynamic interchange between the morphologies; NE tau inclusions (TI-2) underwent morphological changes to TI-3 (speckles) and then to TI-1 (amorphous) inclusions (**Supplementary Figure 4** and **Supplementary Movie 5**, **Figure 3b** and **Supplementary Movie 6**). These data led us to infer that the appearance of ES1 tau inclusions at the NE and cell divisions are mechanistically intertwined in HEK-derived reporter cells, thus contributing to three fluorescent morphologies noted previously (Daude et al., 2020). In the case of ES1 clonal cells, tau inclusions continuously recruit to the NE during the mitotic interphase, appearing as the TI-2 morphology. Loss of NE tau inclusion signals during the cell cycle is consistent with disassembly of the NE and its components as a defining event during metaphase to anaphase transition; this loss of NE inclusion signals was marked by a corresponding increase in TI-3 morphology. During telophase, tau inclusions excluded from NE reassembly fuse together and form large amorphous masses as TI-1 morphology. These processes whereby tau accumulates on the NE and undergoes morphological changes repeat, until a given cell reaches the end of its life span (**Figure 3c, Supplementary Figure 4** and **Supplementary Movie 5**). To determine whether mitotic events are contributing to morphological heterogeneity of tau fluorescent signals, ES1 cells were treated with a cell-cycle arresting reagent, Cyclin-dependent kinase (CDK) 1/2 inhibitor III; this is a cell-permeable inhibitor that targets both CDK1/cyclin B and CDK2/cyclin A and is reported to arrest cells at the G2/M boundary (Jorda et al., 2018). CDK1/2 inhibitor III applied at 10 nM concentration was sufficient to inhibit the proliferation of ES1 cells without overt cytotoxic effects (**Figure 3d**), and concomitantly this same concentration increased the number of cells showing TI-2 morphology (42.6 ± 2.3% compared to control cells 28.7 ± 2.6%; **Figure 3e**), supporting the hypothesis that the tau conformer in the ES1 clonal line has the propensity to bind to NE and undergoes morphological changes as an inevitable consequence of NE disassembly and reassembly.

We then explored whether the NE tau inclusions were associated with cytotoxic effects. Sedimentation analysis revealed that ES1 cells contained mostly insoluble forms of tau, whereas non-seeded reporter cells (4RD-YFP P301L/V377M) had entirely soluble tau (**Figure 4a** and **4b**). While the heterogeneous morphology of NE tau inclusions in ES1 cells persisted for more than 200 days in culture post sub-cloning, the ES1 cells did however show an increase in cleaved lamin B1 (**Figure 4a** and **4b**), a decrease in the cell proliferation (**Figure 4c**) and an increase in lactate dehydrogenase (LDH) activities in the conditioned media compared to the non-seeded reporter cells (**Figure 4d**). These data suggest that an elevated level of cell death might be linked to the presence of NE tau inclusions. Levels of cleaved caspase 3 (Cas-3), Bax dimers and fragmented lamin B1, which are apoptotic cell death markers (Vince et al., 2018; D. Zhang, Beresford, Greenberg, & Lieberman, 2001), were higher in ES1 than in un-transduced reporter cells (**Figure 4e** and **4f**, **Supplementary Figure 5**). Interestingly, time-lapse imaging of live cells revealed apoptosis-like death of multinucleated cells with NE tau inclusions, as characterized by nuclear collapse and formation of apoptotic bodies (**Figure 4g** and **Supplementary Movie 7**).

**Figure 4.**
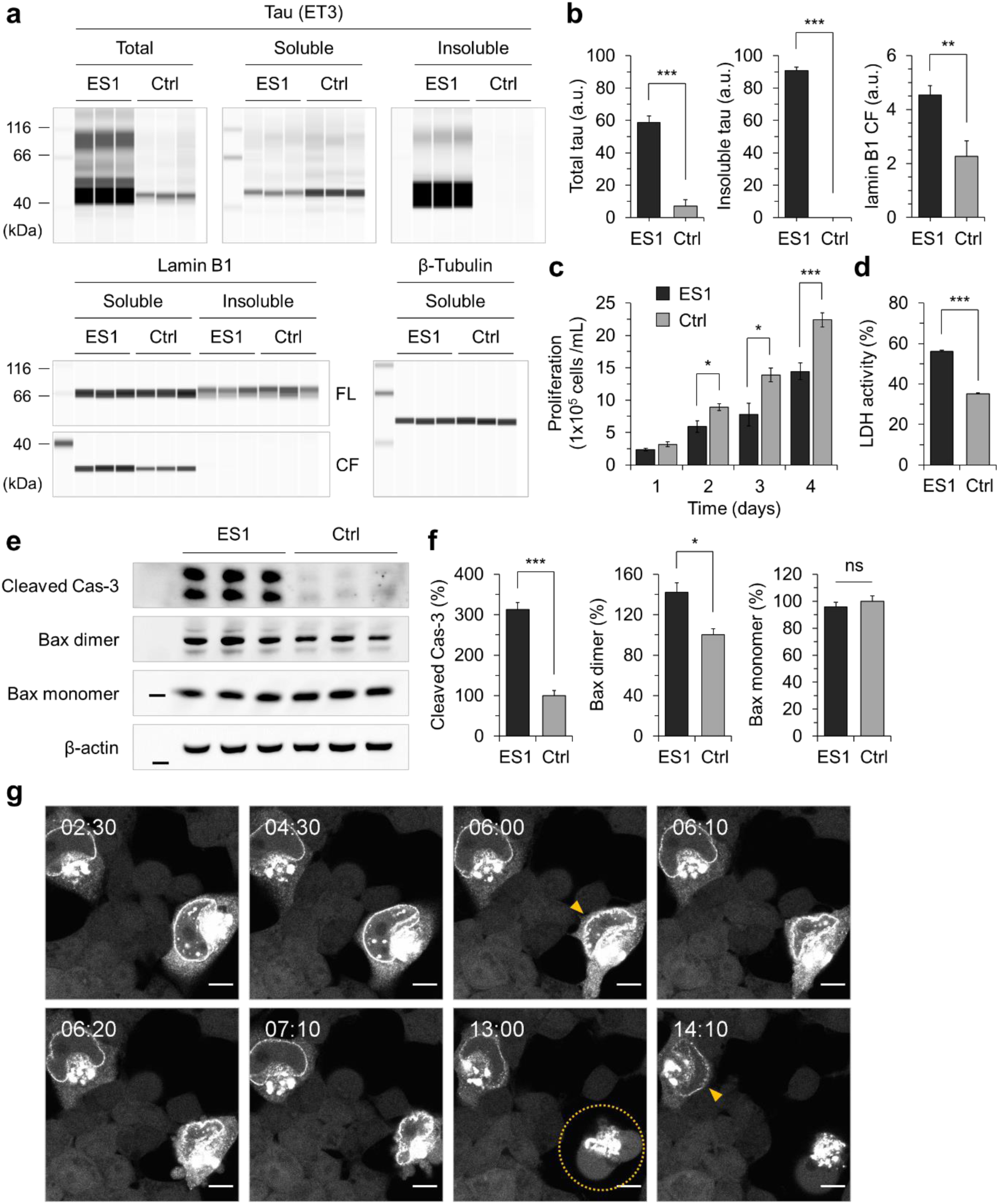
Apoptotic cell death with nuclear envelope tau inclusions. **a.** Sedimentation of Triton X-100 insoluble tau. ES1 and control cells (the reporter cells, ‘Ctrl’) were lysed in PBS-T and sedimented at 100,000xg for 1 hour. The amount of tau and lamin B1 (FL, 66 kDa full-length; CF, 31 kDa cleaved C-terminal fragment) were analyzed using capillary western using anti-tau mAb (ET3) and anti-lamin B1 pAb, respectively (n=3). β-tubulin, a loading control. **b.** Intensity measurement of the capillary western results in (**a**). Intensities were normalized to those of β-tubulin. a.u., arbitrary units. **c.** Proliferation of ES1 cells were determined by counting viable cells at the indicated time points (n=4). **d.** LDH activity in ES1 conditioned media were measured as an indicative of cell death at 3 days post splitting (n=4). **e.** Western blot analysis of apoptosis in ES1 cells. The amount of cleaved caspase 3 and dimerized Bax were analyzed in ES1 cells and control cells (tau reporter cells, Ctrl) (n=3). **f.** Intensity measurement of the western results in (**a**). Intensities were normalized to those of β-actin. **g.** Live cell imaging of tau reporter cells seeded with tau as per Figure 2. Cells with NE tau inclusions underwent apoptotic cell death (yellow circle) followed by nuclear deformation (yellow arrowheads). Cells were imaged every 10 min for 16 hours. Scale bar, 10 μm. Error bars represent SEM. \**p* < 0.05, \*\**p*<0.01 and \*\*\**p* < 0.001 in comparison with the controls.

### Cytotoxicity potential of NE-associated tau inclusions

Building on the above, we considered whether NE accumulation of tau inclusions might interfere with the functionality of NPCs, and thereby cause disrupted NCT, noting that NPCs reside in the NE and mediate bidirectional NCT of molecules essential for cell proliferation and survival (Beck & Hurt, 2017; Strambio-De-Castillia, Niepel, & Rout, 2010). It is reported that under certain conditions of tauopathy, tau binds to NUPs (Eftekharzadeh et al., 2018), which are the main components of the NPCs and embedded in the central lumen of NPCs. We used immunocytochemistry with the anti-NUPs mAb NPC414 (detecting conserved Phe and Gly-rich repeats on NUPs 62, 90 and 152) and an anti-NUP98 pAb to confirm a mis-localization of NUPs in the presence of tau inclusions (**Figure 5a** and **Supplementary Figure 6**). Nuclear deformation and/or bubble-like protrusions on the nuclei were evident in ES1 cells with a large mass of tau inclusions (**Figure 5b**), overlapping some observations in a previous report that pathogenic mutations in tau can cause microtubule-mediated deformation of nuclei, as seen in post mortem analyses of tissues (Paonessa et al., 2019).

**Figure 5.**
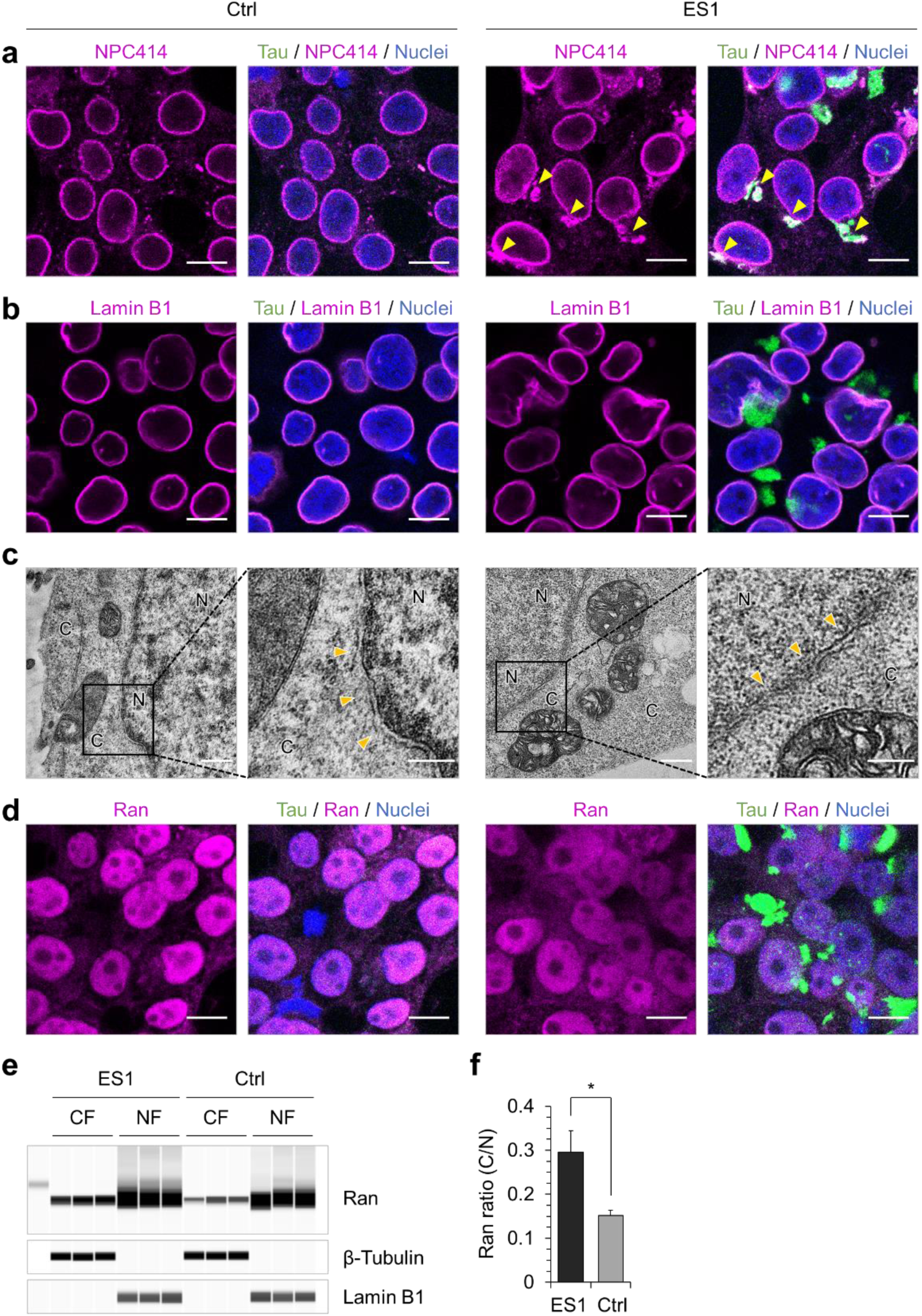
Mis-localization of nucleoporins and disruption of Ran gradient. **a** and **b**. Mis-localization of nucleoporins (NUPs). Tau reporter cells (Ctrl, 4RD-YFP P301L/V337M) and ES1 cells were fixed and permeabilized. NUPs (**a**) and lamin B1 (**b**) were probed and visualized with fluorescent conjugated secondary antibodies. Tau in green; NUPs and lamin B1 in magenta; nuclei were counterstained with DAPI (blue). Localization of nucleoporins with tau were indicated by yellow arrowheads in (**a**). Scale bar, 10 μm. **c**. TEM analysis of ES1 cells showed a disruption of the double membrane architecture of NE in comparison with control cells (Ctrl). Arrowheads indicate NE. C, cytoplasm; N, nucleoplasm. Scale bar, 500 nm and 250 nm in the boxed images. **d** to **f**. Disruption of Ran gradient. **d**. Ran in tau reporter cells (Ctrl) and ES1 cells were probed by immunocytochemistry as described in (**a**) and (**b**). Tau in green; Ran in magenta; nuclei were counterstained with DAPI (blue). **e**. To determine Ran gradient in tau reporter cells and ES1 cells, cytoplasmic and nuclear fractions were separated by differential detergent fractionation and analyzed using capillary western (n=3). **f**. Intensities of the capillary western results in (**d**) were normalized to those of β-tubulin (cytoplasmic fractions) or lamin B1 (nuclear fractions). Ran ratio (C/N), ratios of the cytoplasmic concentration to the nuclear concentration. Scale bar, 10 μm. Error bars represent SEM. \**p* < 0.05 in comparison with the controls.

The NE itself consists of the inner and outer nuclear membranes, which are separated by the perinuclear space (Guttinger et al., 2009; Suntharalingam & Wente, 2003). Transmission electron microscopy (TEM) revealed that the structural integrity of the double-layered NE was ruptured in ES1 cells versus controls (**Figure 5c**). We also investigated Ran, a small GTP-binding nuclear protein involved in the regulation of NCT of RNAs and proteins; Ran shuttles across the NPCs, but is concentrated in the nucleus due to the active delivery mediated by nuclear transport factor-2. This bias in partitioning is known as the Ran gradient (Clarke & Zhang, 2008). Immunocytochemistry and capillary western analysis of cytoplasmic and nuclear fractions indeed confirmed a decline of the Ran gradient in ES1 cells compared to un-transduced parental reporter cells (**Figure 5d** to **5f**). These data support a view that NE tau inclusions in ES1 cells trigger the separation of NUPs from NPCs, disrupt molecular trafficking across the nuclear envelope, and thereby contribute to cellular dysfunction that may then trigger programmed cell death pathways.

### Dynamic analyses of nuclear-cytoplasmic transport

For dynamic analyses of NCT, tau reporter cells (4RD-YFP P301L/V377M) and the ES1 subline were transiently transfected with an additional type of reporter construct, a plasmid encoding two proteins indicating the status of nuclear-cytoplasmic compartmentalization (NCC) events. The NCC reporter plasmid (pLVX-EF1alpha-2xGFP:NES-IRES-2xRFP:NLS) has been used by others (Eftekharzadeh et al., 2018; Mertens et al., 2015; Paonessa et al., 2019); it encodes a nuclear export signal (NES) fused to green fluorescent protein (GFP) and a nuclear localization signal (NLS) fused to red fluorescent protein (RFP) under the control of a human elongation factor-1α (EF-1α) promoter and with an internal ribosome entry sequence (IRES) being located between the two open reading frames (**Figure 6a**) (Mertens et al., 2015). Corresponding tau reporter cells exhibited a segregated arrangement of the fluorescent signals; GFP localized in cytoplasm and RFP localized in nuclei (**Figure 6b** to **6d**). Although intense YFP signals of tau inclusion in ES1 cells hindered the analysis of NES-GFP compartmentalization, the increased levels of local RFP signals in both nuclei and cytoplasm indicates an impairment of NCC (**Figure 6b**). The impaired NCC in ES1 cells became evident upon quantifying intensities of the pixels along a chord (dotted line with arrow) placed across the cell bodies and nuclei of transfected cells (**Figure 6c** to **6f**).

**Figure 6.**
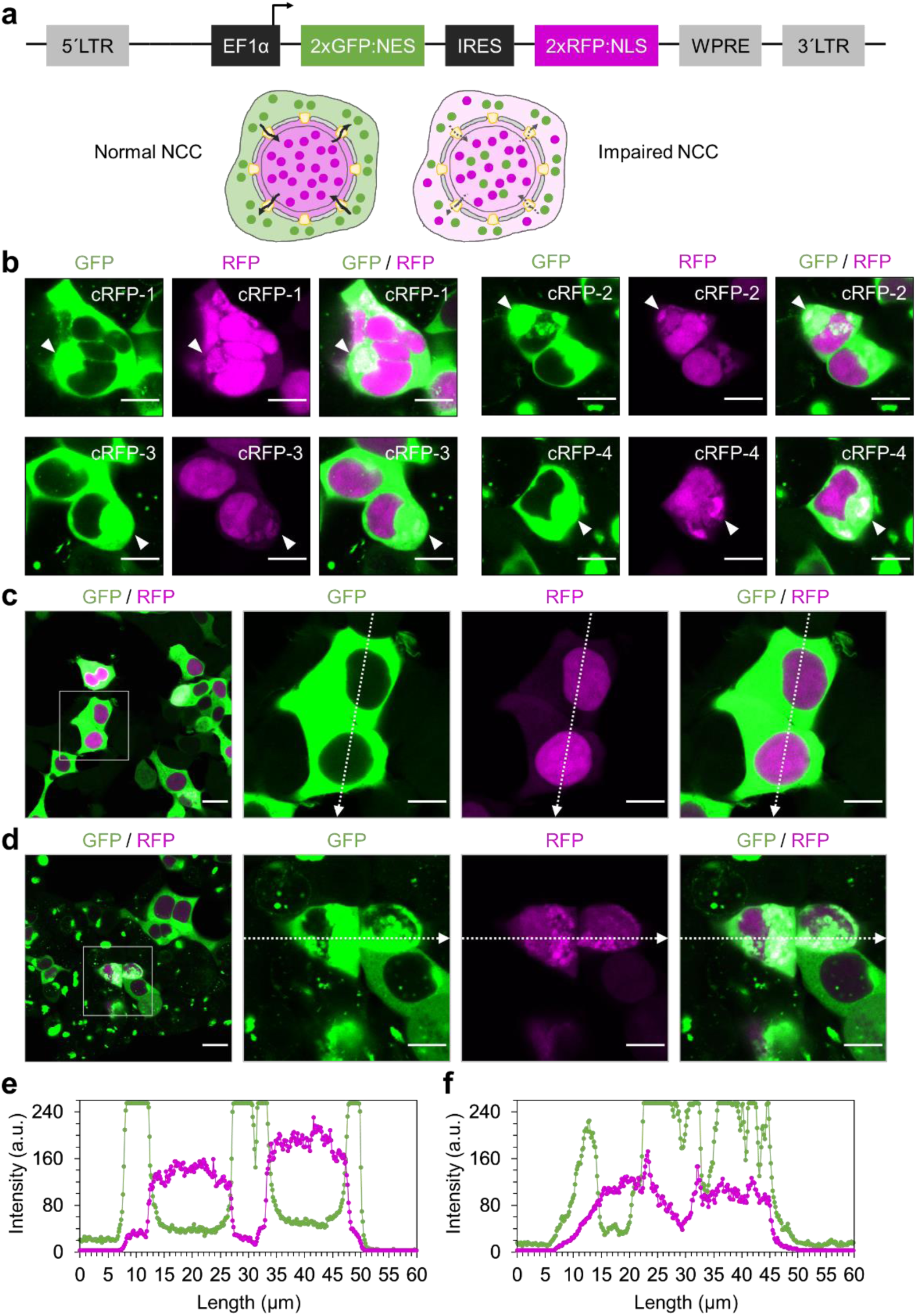
Defects in nuclear-cytoplasmic compartmentalization. **a**. Nuclear-cytoplasmic compartmentalization (NCC) reporter construct. NCC reporter encodes GFP and RFP (colored in magenta) fused with a nuclear export signal (NES) and a nuclear localization signal (NLS), respectively. Schematic illustration shows co-localization of GFP and RFP, indicating NCC defects under an impaired NCC condition. **b**. Cytoplasmic localization of NLS-RFP in ES1 cells was indicated by arrowheads. cRFP, cytoplasmic RFP. Scale bar, 10 μm. **c** and **d**. Tau reporter cells (**b**) and ES1 cells (**c**) were transiently transfected with NCC reporter construct and imaged after 24 hours. Scale bar, 20 μm and 10 μm in the boxed images. **e** and **f**. Intensities of green and red fluorescence signals (colored in magenta) in the reporter cells (**c**) and ES1 cells (**d**) were measured along the arrows with a length of 60 μm.

ES1 transfected cells harboring the pLVX-EF1alpha-2xGFP:NES-IRES-2xRFP:NLS plasmid showing the anticipated segregated pattern of the fluorescent signals were then subjected to a “fluorescence recovery after photobleaching” (FRAP) analysis (**Figure 7a**). Since the emission spectra of GFP encoded in the NCC reporter and YFP fluorophore fused to the tau repeat domain are overlapped, we restricted ourselves to the use of RFP signals for FRAP analyses. RFP signals in the entire images were photobleached, such that nuclear RFP signals appearing *de novo* in the field of view must derive from newly-synthesized molecules. Five initial time-lapse images were taken as points of reference, with subsequent recovery of signal in the RFP channel measured every 10 min thereafter for 6 hours (**Figure 7b**, **Supplementary Movie 8** and **Movie 9**). Nuclear RFP signals in tau reporter cells recovered within an hour after photobleaching up to 16.4 ± 2.2% and reached to 32.0 ± 7.0%; the ES1 cells on the other hand showed a slower recovery, with only 4.5 ± 1.0 % signal at one hour and a final attained value of 9.6 ± 1.9% (**Figure 7c** and **7d**; all figures compared to an average of the reference RFP intensities). These observations are consistent with defects in the selective NE permeability seen in induced pluripotent stem cells (iPSCs)-derived neurons with IVS10+16 and P301L *MAPT* mutations (Paonessa et al., 2019), and in primary neurons treated with high molecular weight (HMW) AD brain fractions containing tau (Eftekharzadeh et al., 2018).

**Figure 7.**
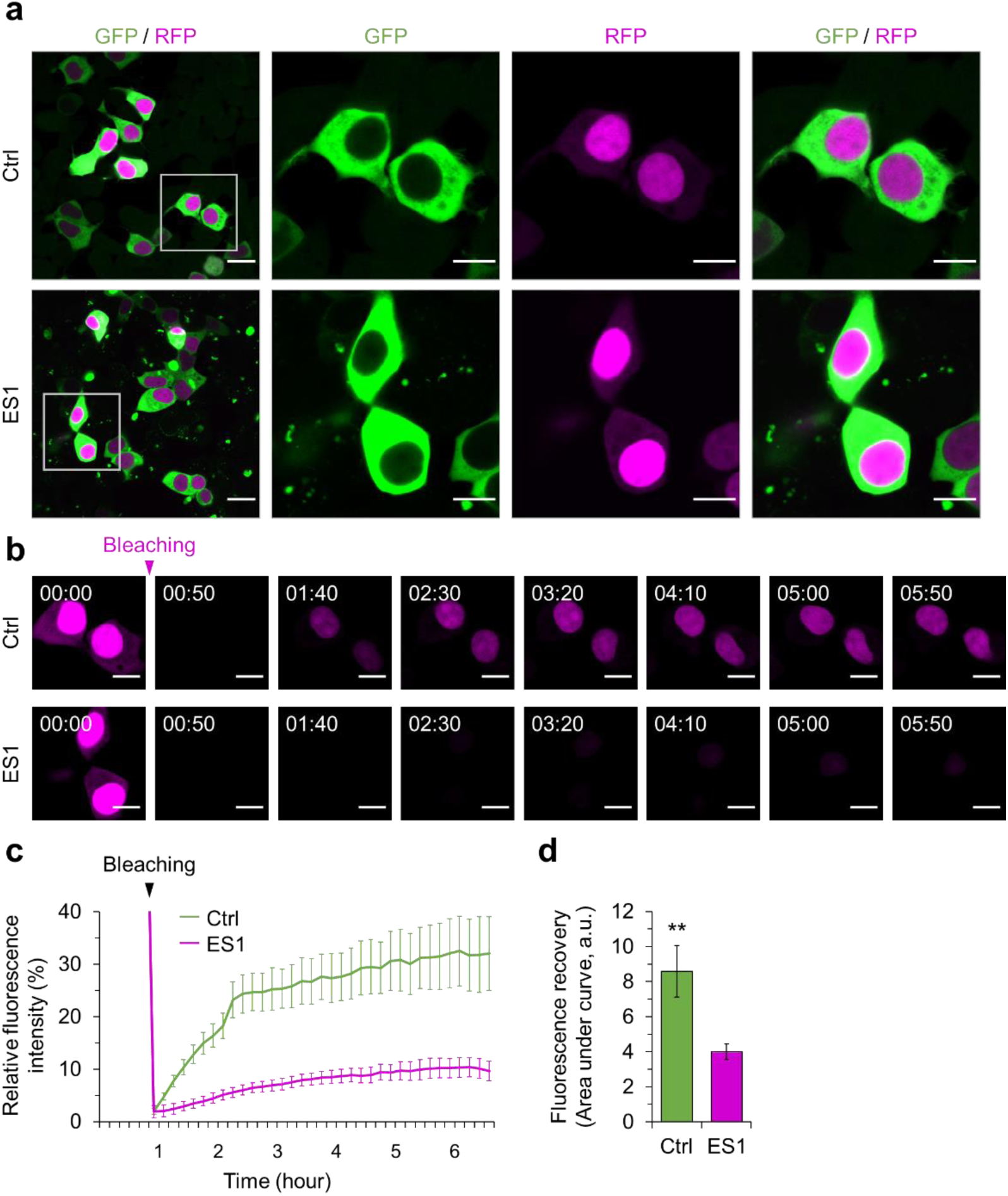
Disruption of nuclear-cytoplasmic transport. **a**. Tau reporter cells (Ctrl) and ES1 cells transfected with NCC reporter construct were imaged after 24 hours. Cells showing a normal nuclear-cytoplasmic compartmentalization were subjected to fluorescence recovery after photobleaching (FRAP) analysis. Scale bar, 20 μm and 10 μm in the boxed images. **b**. Live cell imaging of the FRAP analysis. Red fluorescence signals (colored in magenta) were completely photobleached and then images were obtained every 10 min for 7 hours. The magenta arrowhead indicates the time when photobleaching was applied. Scale bar, 10 μm. **c**. Realtime measurements of fluorescence recovery of nuclear red signals in tau reporter cells and ES1 cells (n=8). **d**. The data were presented as accumulated signals under the average curves in (**c**). Error bars represent SEM. \*\**p* < 0.01 in comparison with the controls (Ctrl).

### A demixed liquid state of oligomeric tau on nuclear envelope

We next sought evidence for the *in vivo* formation of a liquid state of tau. In tau reporter cells (4RD-YFP P301L/V377M) seeded with brain homogenate of CSA Type 2 conformers, dispersed tau-YFP signals were sequestered to tau inclusion with various morphologies (**Supplementary Figure 7a**). Quantification of image pixels demonstrated condensation of tau occurred by a seeding reaction, in which signal intensities were concentrated on NE tau inclusions (**Figure 8a** and **8b**, **Supplementary Figure 7b** and **7c**). The average intensity in the cell bodies of tau reporter cells was 23.7 ± 0.2 arbitrary units (a.u.), while the seeded reporter cells with TI-2 morphology showed 14.1±0.1 a.u. (**Figure 8b**). In stable ES1 subline, photobleached NE tau inclusions were rapidly recovered (within 15 min) in FRAP analysis. Notably, different focal plane images revealed that tau inclusions showed liquid droplet-like movements and fused together to increase in size (**Figure 8c** and **8d**, **Supplementary Movie 10**). While relatively large inclusions such as juxtanuclear inclusions had little ability to recover (**Figure 8e**, **Supplementary Movie 11**). These properties meet common criteria for defining a phase-separated structure under live cell conditions, namely spherical morphology, fusion events and recovery from photobleaching (Alberti et al., 2019).

**Figure 8.**
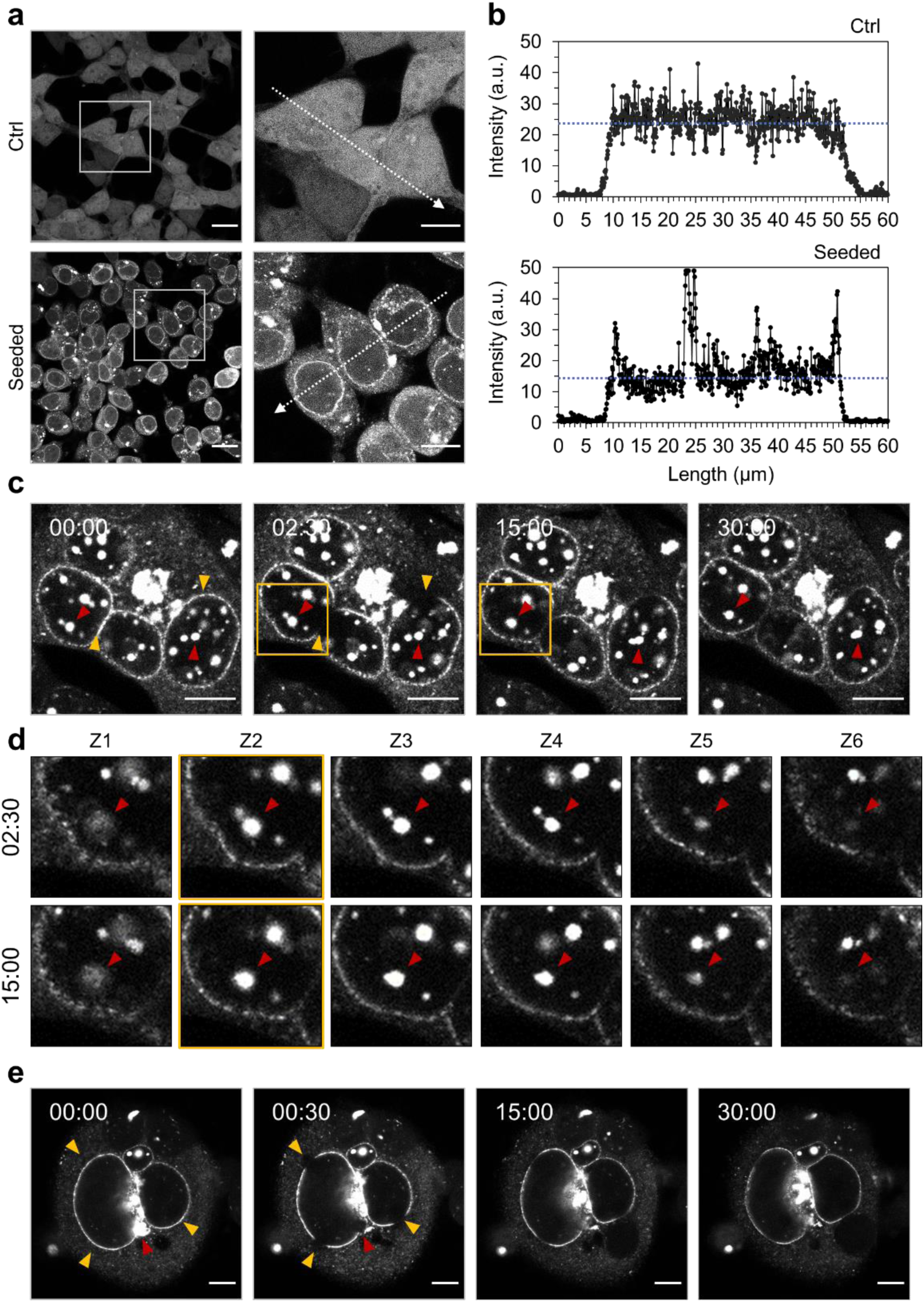
Condensation of demixed tau droplets on nuclear envelope. **a** and **b**. Condensation of tau protein occurred with tau seeding. Tau reporter cells (4RD-YFP P301L/V377M, Ctrl) were seeded with tau and imaged as per Figure 2 (**a**). Intensities of tau signals were measured along the arrows with a length of 60 μm (**b**). a.u., arbitrary units. **c**. Fluorescence recovery after photobleaching (FRAP) analysis of NE tau and fusion of droplet-like tau inclusions. NE tau signals were photobleached (yellow arrowheads) at the indicated time point and then images were obtained every 30 sec for 30 min. Droplet-like tau inclusions fused together (red arrowheads). **d**. Different focal plane images of the boxed areas in (**c**). Z1 to Z6 are depths of field from bottom to top with 1 μm intervals. Arrowheads indicate tau inclusions fused into one droplet. **e**. FRAP analysis of NE tau (yellow arrowheads) and amorphous tau inclusion (a red arrowhead) in multinucleated cells was performed as per (**c**). Scale bar, 10 μm.

To further confirm LLPS of tau in response to seeding by exogenous misfolded tau, ES1 cells exhibiting a heterogeneous repertoire of tau inclusion phenotypes were stained with thioflavin S (ThS), which shows an increase in the emission of a fluorescent signals upon by binding to fibrillar assemblies (Wegmann et al., 2018; Xu, Martini-Stoica, & Zheng, 2016). ThS staining readily visualized amorphous and juxtanuclear inclusions in these analyses (TI-1), but - crucially - not NE inclusions (TI-2), nor small speckles (TI-3) (**Figure 9a**). Intriguingly, particle size plotted against fluorescent signal for the YFP and ThS double-positive inclusions revealed that YFP-associated area under the curve was always wider than ThS area (**Figure 9b**), thus indicating that aggregated tau fibrils (ThS-positive) have a surrounding milieu of condensed tau existing in a liquid state. The inconsistency in the ratio of ThS to YFP may indicate a potential involvement of molecular interactions with other polymers in liquid-solid phase transition under live cell conditions. Particle size distribution in these experiments demonstrated that tau inclusions less than 1 μm^2^ were dominant in the YFP-only population, while the number of YFP and ThS double-positive particles instead peaked in the size range 1 to 3 μm^2^ (**Figure 9c**). Taken together, these data indicate that dispersed soluble form of cellular tau condensed and underwent LLPS under conditions of tau seeding. Importantly, the primary nucleation of tau fibrils, which has been inferred from observations under conditions of molecular crowding (Ambadipudi et al., 2017; Fichou et al., 2018; Wegmann et al., 2018; X. Zhang et al., 2017), was demonstrated under here within living cells, with condensed tau droplets ranging in size from 1 to 3 μm^2^ in diameter (**Figure 9d**) and with the larger droplets capable of producing fibrillar tau in their interior.

**Figure 9.**
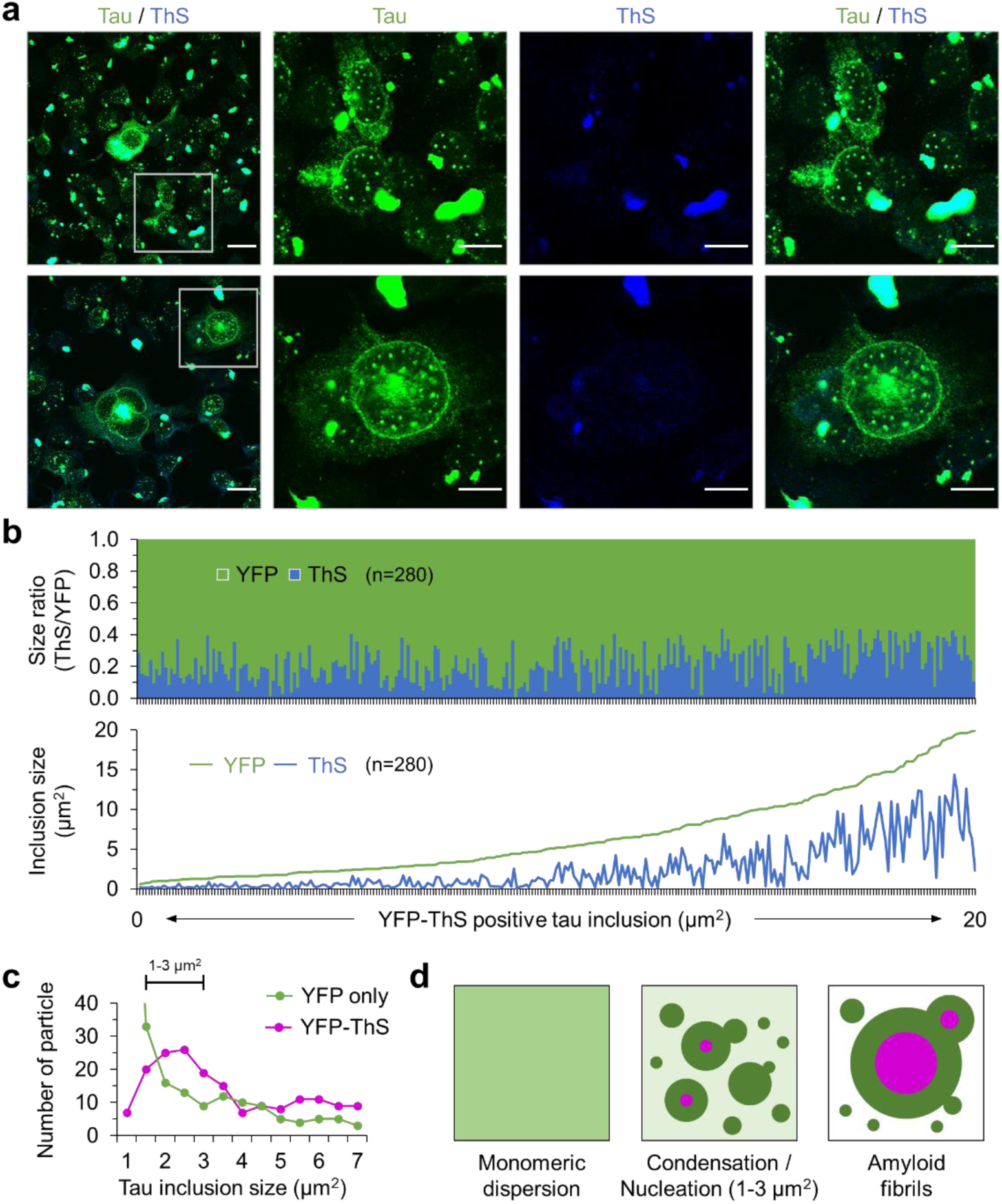
Nucleation of amyloid fibrils in tau droplets. **a**. Amyloid fibril formation in ES1 cells. The cells were stained with Thioflavin S (ThS), which increased its fluorescence upon binding to β-sheet structure in aggregated amyloid fibrils. Scale bar, 20 μm and 10 μm in the boxed images. **b**. Area measurements of YFP-ThS double positive tau inclusions observed in (**a**). Total 280 particles were analyzed and presented as a ratio of ThS to YFP (top) and the area of ThS and YFP within each inclusion (bottom). **c**. Size distributions of YFP only positive (n=176) and YFP-ThS double positive (n=263) tau inclusions. The majority of YFP only positive tau inclusions were smaller than 1 μm^2^ (n=139). **d**. Schematic illustration shows a condensation of monomeric dispersed tau protein into demixed liquid droplets (dark green). Primary nucleation of tau (magenta) occurs in droplets with a size of 1-3 μm^2^. Tau fibrils grow further by recruiting condensed tau droplets and droplets including small fibrils.

## Discussion

Perturbations of NCT have been observed in neurodegenerative disorders with a number of protein aggregates and cytoplasmic assemblies, including artificial β-sheet deposits, Huntingtin inclusions, α-synuclein aggregates, SGs, and C9ORF72 G_4_C_2_ RNA assemblies (Grima et al., 2017; Jiang et al., 2016; Jovicic et al., 2015; Woerner et al., 2016; K. Zhang et al., 2018; K. Zhang et al., 2015). For tauopathies, disruption of NCC with high molecular weight (HMW) tau species derived from AD brain and *MAPT* mutation-mediated NE deformation have been observed in cortical primary neurons and in neurons derived from induced pluripotent stem cells (iPSCs), respectively (Eftekharzadeh et al., 2018; Paonessa et al., 2019). We have reported NE tau inclusions as the predominant morphology in 4RD-YFP P301L/V377M reporter cells seeded with misfolded tau conformers (CSA Type 2) found in the brains of some aged TgTau^P301L^ mice or cortical samples from certain FTLD-MAPT patients given a clinical diagnosis of bvFTD (Daude et al., 2020; Eskandari-Sedighi et al., 2017). Here we have extended this observation by establishing a stable subclone, ES1, using CSA Type 2 seeds. ES1 clonal cells partly resemble DS9 clonal cell line, that propagates synthetic strains derived from recombinant tau (Sanders et al., 2014; Sharma, Thomas, Woodard, Kashmer, & Diamond, 2018), with regards to Triton X-100 insoluble tau and a 12 kDa product (as the ‘core’ of the amyloid) after pronase E digestion, but, rather than the speckle-shaped inclusions of DS9 cells, they harbor NE and heterogeneous fluorescent tau inclusion morphologies. Interestingly, DS10, which is the other clonal line propagating the synthetic strains, created multiple stably sub-strains easily discerned by different tau inclusion morphologies (Sharma et al., 2018), whereas ES1 clone derived from brain materials with CAS Type 2 profile produced a single population of six clones, all identical to ES1 in tau inclusion morphology (**Figure 3a** and **Supplementary Figure 3a**).

Fluorescent tau deposits in immortalized 4RD-YFP reporter cells (4RD-YFP P301L/V377M) or GFP-0N4R reporter cells (Dox:GFP-0N4R P301L) exposed to CSA Type 2 seeds exhibited heterogeneous morphologies, but with NE inclusions being prominent amongst these. This phenotypic property of tau inclusions persisted after the single cell cloning (ES1), while, compared with non-seeded control cells, the proliferation rate was lower with loss of cells by apoptosis becoming evident. This decline in cell viability following tau seeding activity mirrors previous observations for certain types of tau seeds (Kaufman et al., 2016; Sanders et al., 2014). The NE tau inclusions and decrease in cell viability led us to deduce that interference with NCT is the pathogenic mechanism of tauopathies at the cellular level. This hypothesis is strongly supported by further findings on ES1 cells including mis-localization of NUPs into tau inclusions, alteration in the ratio of nuclear to cytoplasmic concentration for the Ran protein and a decline in NCC. An alternative explanation is that separation of NUPs from NPCs occurs earlier, with mis-localized NUPs then binding to intrinsically disordered tau and appearing as NE tau inclusions, but this theory of indirect action then begs the question of the proximal cause of NUPs dissociation.

Phase transition into demixed liquid state of tau has been reported mainly using *in vitro* cell-free systems with purified recombinant proteins under conditions of molecular crowding (Ambadipudi et al., 2017; Boyko et al., 2019; Hernandez-Vega et al., 2017; Majumdar, Dogra, Maity, & Mukhopadhyay, 2019; Singh et al., 2020; Vega et al., 2019; Wegmann et al., 2018; X. Zhang et al., 2017). Intrinsic and documented aspects of tau biology include natively disordered structure, inhomogeneous charge distribution, variable patterns of physiological and pathological phosphorylation, pathogenic mutations, and alternative splicing sites producing six different isoforms, any and/or all of which might lead to increased acquisition of LLPS (Ambadipudi et al., 2017; Boyko et al., 2019; Wegmann et al., 2018; X. Zhang et al., 2017). In cultured cells, a GFP-tagged version of the longest isoform of wild-type tau (GFP-tau441) formed droplet-like accumulations in transiently transfected mouse primary cortical neurons and N2a neuroblastoma cells with high expression levels (Wegmann et al., 2018). Increased local concentration of aggregation prone proteins, such as pathogenic TDP43, hnRNPA1 and FUS, has been considered to enhance protein interactions causing LLPS (Harrison & Shorter, 2017; Molliex et al., 2015; Murakami et al., 2015; Shin & Brangwynne, 2017). In this study, a tau RD domain/YFP fusion protein with pathogenic mutations on the repeat domain, P301L and P301L/V337M, is stably dispersed throughout the cytoplasm and the entire cell body, without forming protein clusters. The final concentration of total tau used to seed the reporter cells, including soluble and insoluble forms in the presence of sarkosyl, was only 20 ng/mL based on the estimation using CDI (Daude et al., 2020) whereas signal for total tau in ES1 cells was approximately 8 times greater than that for in non-seeded controls, as analyzed by capillary western (**Figure 4a** and **4b**). These data strongly suggest that the tau condensation on NE and inclusion formation occurred in response to the pathogenic tau seeding rather than a hypothetical redistribution effect causing a locally increased tau concentration. Moreover, the NE tau inclusions and small droplets were found to behave as an oligomeric liquid phase (as determined by a combination of FRAP, live cell imaging analysis, and amyloid fibril staining with ThS described below), with implications for the cell biology of disease pathogenesis (**Figure 10**).

**Figure 10.**
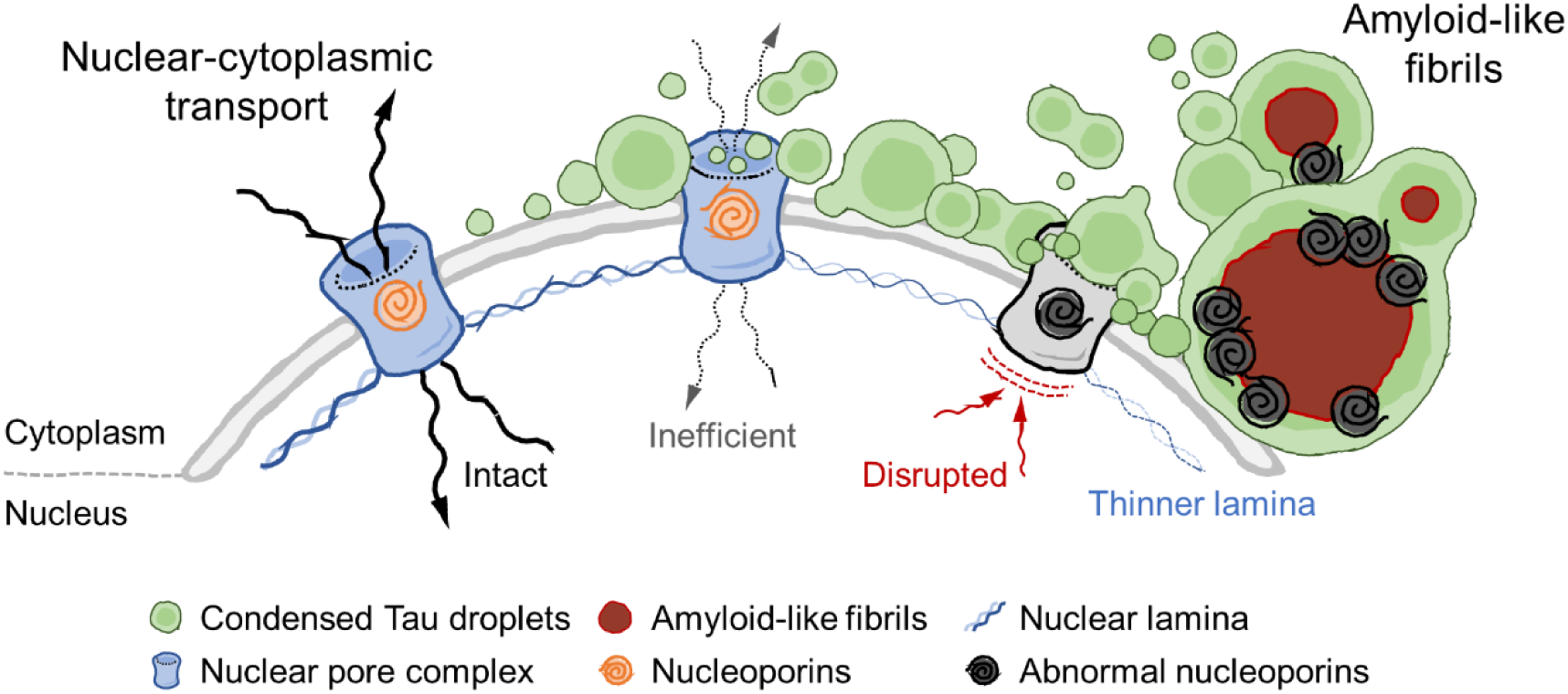
Graphic summary. Condensed tau droplets perturb the nuclear envelope. Under pathogenic conditions (e.g. pathogenic tau seeding), dispersed tau condense as liquid droplets and recruit to the nuclear envelope, resulting in a decline in NCT. Continuing tau LLPS and condensation cause mis-localization of NUPs and completely disrupt molecular trafficking between the nucleus and cytoplasm. A liquid-solid phase transition (i.e. primary nucleation) occurs in the core of the liquid droplets and grows as amyloid-like fibrils.

Concerning the pathways leading to cell death, soluble oligomeric, but not monomeric nor fibrillar, forms of tau have long been considered to be cytotoxic due to their ability to internalize into recipient cells and recruit monomeric tau into filamentous inclusions (tau seeding activity) (Flach et al., 2012; Frost, Jacks, & Diamond, 2009; L. Guo & Lee, 2011; Lasagna-Reeves, Castillo-Carranza, Guerrero-Muoz, Jackson, & Kayed, 2010; Lasagna-Reeves et al., 2011; Patterson et al., 2011; Rauch et al., 2020). In addition and more recently, several lines of evidence suggest that tau uptake and aggregation are not sufficient *per se* to cause immediate neuronal cell death (Ait-Bouziad et al., 2017; Takeda et al., 2015). Moreover, the presence of neurofibrillary tangles (NFTs) does not inevitably lead to neuronal and network dysfunction *in vivo* (Kuchibhotla et al., 2014). Taken together, these data imply a toxic intermediate which may be independent of the processes for internalization and propagation of intracellular tau aggregates. Experiments described here show that a liquid state condensed tau resulting from the introduction of pathogenic seeds, at least the tau conformer prominent in CSA Type 2 brain tissues, is recruited to the NE and triggers a disruption of NCT. In turn, the response of the compromised cells is to initiate a regulated cell death pathway, this pathway having the hallmarks of apoptosis in the HEK-derived ES1 cells, as shown by production of cleaved caspase 3, accumulation of Bax dimers, production of cleaved product of lamin B1 and light microscopic observation of apoptotic bodies (**Figure 4**).

The scheme for pathogenesis outlined above aligns with neuropathological data for FTLD-MAPT and new insights into how tau conformers in the brain are not homogeneous but occur in ensembles (Daude et al., 2020). For tau staining, juxtanuclear signals seen in TI-1 and TI-2 morphologies in acute protein transduction and ES1 cells produced by seeding with material assigned as a CSA Type 2 conformer signature (**Figure 2a**, **Figure 3e** and **Supplementary Figure 1**) have a parallel in terms of focal tau immunostaining of some DG neurons of TgTau^P301L^ mice (**Figure 1q**) and in “mini Pick-like bodies” of FTLD-MAPT-P301L cases (Borrego-Ecija et al., 2017). For lamin B1 alterations reflecting changes in nuclear function and architecture as well as secondary alterations induced by apoptosis-associated caspases action, irregular nuclear margins are seen in FTLD-MAPT-P301L carriers and in aged TgTau^P301L^ mice (**Figure 1f** and **1m**), as well as focal diminutions of signal intensity (**Figure 1d** and **Figure 1n**); direct physical interactions between tau and lamin B1 however are not supported, given double-staining results in DG neurons and analyses of Triton X-100 insoluble ES1 cell fractions. Cytoplasmic lamin B1 puncta in human brains (**Figure 1g** and **1h**) may reflect apoptotic bodies from partitioning of cellular contents. Conversely, nuclear clefts detected with lamin B1 antibody were also present in other neurodegenerative diseases (**Table 1**) and in aged non-Tg mice (**Figure 1k** and **1l**) and were not observed in protein transduced HEK cells and stably-transduced ES1 cells; they thus reflect age-dependent changes but could nonetheless comprise a comorbidity to exacerbate toxic effects of tau accumulation.

In sum, our data support a central toxic mechanism where tau conformer ensembles that accumulate in bvFTD, especially CSA Type 2, give rise to demixed oligomeric tau forms that initiate neuronal death by binding to the NE. In transduced reporter cells these forms are associated with fluorescent tau inclusion (TI) morphologies TI-1 to TI-3. An interlocking perspective, participation in a downstream pathogenic pathway, may apply to a fourth morphology scored as cytoplasmic threads (TI-4). Thus, the TgTau^P301L^ mouse brains yielding most TI-4 morphologies in protein transduction experiments have florid pathology with AT8 phospho-tau antibody that includes numerous tangle-like deposits. Greater than 96% of the structures scored by EM analysis of these brains are straight fibrils and the corresponding tau conformer profile (CSA Type 4) resembles that of recombinant tau fibrils (Daude et al., 2020; Eskandari-Sedighi et al., 2017). These data for TI-4 can be reconciled with the views that a) tangles and filamentous tau are less toxic than oligomeric tau forms (Flach et al., 2012; Frost et al., 2009; J. L. Guo & Lee, 2011; Kuchibhotla et al., 2014; Lasagna-Reeves et al., 2010; Lasagna-Reeves et al., 2011; Patterson et al., 2011; Rauch et al., 2020) and that b) a later-stage *in vivo* event is a liquid to solid phase transition from LLPS tau that nucleates intracellular tau fibrils ultimately giving rise to fibrillar tau seen by light microscopy in FTLD-MAPT cases. In other words, once LLPS occurs it may serve as a nursery for the fibrillar tau forms seen at disease endpoint (Kim et al., 2013; Lee et al., 2016; Mann et al., 2019). The hierarchy of pathogenic events deduced from our results would seem to have two implications. First, FTLD-MAPT is a 4R-tauopathy that necessarily encompasses tau accumulation in astrocytes and oligodendroglia representing diverse pathologies in tauopathy (Gotz et al., 2019), as well as in neurons; how LLPS phenomena might operate in these other cell lineages and under conditions of neuroinflammation is wide open and worthy of investigation. Second, the transition to the ensemble of tau conformers defined by a CSA Type 2 profile from a “cloud” of conformers in a prodromal state (Daude et al., 2020) would appear to be a crucial point for disease intervention.

## Material and methods

### Ethics statement

Ethical review at the University of Alberta was performed by the Research Ethics Management Office, protocols AUP00000356 and Pro00079472. All other procedures were performed under protocols approved by the Institutional Review Board at the IDIBAPS brain bank (Barcelona, Spain). In all cases, written informed consent for research was obtained from patients or legal guardians and the material used had appropriate ethical approval for use in this project.

### Brain tissues of patients and transgenic mice and immunohistochemistry

FTLD-MAPT-P301L patients of both sexes were as described previously (Borrego-Ecija et al., 2017) and as per **Table 1**. Clinical features of the patients were assessed as per contemporaneous criteria for diagnosis (Gorno-Tempini et al., 2011; Rascovsky et al., 2011). Control brain samples were obtained from patients who died from non-neurological diseases; diagnostic neuropathology and retrospective chart reviews were carried out for all subjects, with particular attention to ruling out other age-related neurodegenerative diseases as previously described (Daude et al., 2020). TgTau^P301L^ mice samples were obtained as described previously (Daude et al., 2020; Eskandari-Sedighi et al., 2017; Murakami et al., 2006). All animal experiments were performed in accordance with local and Canadian Council on Animal Care ethics guidelines.

Brain tissues from patients and transgenic mice were processed for histologic and immunohistochemical purposes as described previously (Eskandari-Sedighi et al., 2017). Briefly, each specimen was fixed in neutral buffered 10% formalin and paraffin-embedded. Six μm sagittal sections were rehydrated and endogenous peroxidase activity was blocked by treatment with 3% hydrogen peroxide for 6 min. The sections were then incubated with primary antibodies at 4°C overnight: anti-phospho-tau mAb, AT8 (1:200, MN1020, Thermo Fisher); anti-Lamin B1 pAb (1:200, ab16048, Abcam). The target molecules were visualized with horseradish peroxidase using the DAKO ARK kit according to the manufacturer’s instruction or with fluorescent-conjugated secondary antibodies: goat anti-mouse IgG (H+L) with Alexa Fluor 594 (Invitrogen, A32742); goat anti-rabbit IgG (H+L) with Alexa Fluor 488 (Invitrogen, A32731). Nuclei were counterstained using Mayer’s hematoxylin or Hoechst 33342 (Invitrogen, H1399), dehydrated and cover-slipped with permanent mounting medium. The section images were acquired with NanoZoomer 2.0-RS digital slide scanner (Hamamatsu) and analyzed using NDP.view2 (Hamamatsu) and Image J software (https://imagej.nih.gov/ij/). Assessment of lamin B immunohistochemistry in the dentate gyrus (DG) of FTLD-MAPT-P301L cases was performed in a semiquantitative way by two observers at a multiheaded microscope at 40x magnification. For clefts, cytoplasmic staining and discontinuous staining intensity: 0.5 = rare, 1-5 neurons affected in one field; 1 = mild, 1-5 neurons affected per field in more than one field; 2 = moderate, 6-15 neurons per field: moderate; 3= severe, >15 neurons affected per field. A similar scheme was used for angular nuclear margins: 0.5 = rare, 1-5 neurons affected in one field; 1 = mild, 1-5 neurons affected per field in more than one field; 2 = moderate, 6-15 neurons per field or >15 affected per field but little angled; 3= severe, >15 neurons affected per field and highly angulated. Tau pathologies were scored as described previously (Daude et al., 2020; Eskandari-Sedighi et al., 2017).

### Cells and cell culture

A monoclonal HEK293 cell line stably expressing human tau repeat domain (4RD) with aggregation prone mutations (P301L/V377M) fused to YFP (4RD-YFP P301L/V377M) (Sanders et al., 2014) were maintained at 37°C with 5% CO_2_ in the culture media; Dulbecco’s modified Eagle’s Medium (DMEM, 11995-065, Gibco) with high glucose (4.5 g/L) and 2 mM glutamine (Gibco), supplemented with 10% fetal bovine serum (FBS, HyClone) and Penicillin (10 units/mL)-Streptomycin (10 μg/mL) (Gibco). To induce cell-cycle arrest, cells were treated with CDK1/2 inhibitor III (CAS 443798-55-8, Calbiochem) (Jorda et al., 2018) for 24 hours at 10 nM concentration. To generate doxycycline-inducible GFP-0N4R tau reporter line, an enhanced GFP and human WT 0N4R tau sequences were inserted between the *Bam*HI and *Xho*I restriction sites on the pcDNA5/FRT/TO plasmid (Invitrogen). A short linker sequence (ATCGATGCA) was incorporated between the eGFP coding sequence (CDS) and 0N4R tau CDS within the construct. Site directed mutagenesis was performed on the resulting plasmid to generate the P301L mutation in the tau CDS (pcDNA5/FRT/TO/GFP-0N4R P301L). The final plasmid and the Flp recombinase vector (pOG44 plasmid, Invitrogen) were packaged with Lipofectamine2000 (Thermo Fisher) and transfected into the Flp-In T-Rex-293 cell line (Invitrogen) according to manufacturer guidelines. Hygromycin B (Thermo Fisher) was used to select stable integrants which were propagated to generate the final cell line (Dox:GFP-0N4R P301L). To induce expression of GFP-0N4R P301L, doxycycline is added to the culture media at a final concentration of 10 μg/ml.

### Tau cell seeding assay

The reporter cells were seeded as previously described (Daude et al., 2020; Eskandari-Sedighi et al., 2017). Briefly, tau reporter cells were plated at 1×10^6^ cells/well of a 12-well culture plates and, on the next day, seeded with liposome-protein complexes derived from brain homogenate of TgTau^P301L^ ill with signs of neurological disease. Two μL of brain homogenate (5-8 mg/mL protein solution was adjusted by total tau content to 8 μg/mL based on the estimation of conformation-dependent immunoassay) (Daude et al., 2020) were combined with the same volume of Lipofectamine 3000 (L3000-015, Thermo Fisher Scientific) and added to the wells. The cells were then incubated for 6 hours at 37°C and the media containing the liposome-protein complex were replaced with fresh culture media.

### Single cell cloning by limiting dilution

The cells were resuspended and counted using the automated cell counter, Countess (Invitrogen) (see Cell viability assay). Two hundred μL of the cell suspensions with concentration of 3 cells/mL were added to each well of 96-well culture plates. Single cell clones in each well were inspected after 4 days and then at two days intervals. The cell clones ensured as only one center of growth were subcultured and frozen in liquid nitrogen until use.

### Cell viability

Cells were resuspended by trypsinization and stained with the same volume of trypan blue (Invitrogen). The samples were loaded into the chamber ports on one side of the Countess cell counting chamber slide (Invitrogen). Viable and dead cells were counted using the automated cell counter, Countess (Invitrogen). Viability was expressed as a percentage of live cells to total cells counted. Cell viability was also determined based on lactate dehydrogenase (LDH) activity in conditioned culture media using a commercial kit (G1780, Promega) following the manufacturer’s instructions. Culture supernatants were collected and incubated with tetrazolium salt, as the substrate, for 30 min at room temperature. The red formazan products of the enzymatic reaction were quantified using a microtiter plate reader (μQuant, Bio-Tek) at wavelength of 490 nm. The LDH activities were expressed as a percentage to the control conditioned media.

### Immunocytochemistry and live cell imaging

Cells were plated on poly-D-lysine (Sigma) and laminin (Sigma) double coated microscope cover glasses (Thermo Fisher Scientific). For immunocytochemistry, cells were fixed in paraformaldehyde (4%, pH 7.4, Electron Microscopy Sciences) for 15 min and optionally permeabilized with PBS containing Triton X-100 (0.1%). The fixed cells were blocked with 1% BSA in PBST (PBS with 0.1% Tween 20) for 30 min and probed with mAb or pAb at 4°C overnight: anti-NPC proteins mAb (1:2,000, ab24609, abcam); anti-NUP98 pAb (1:2,000, NBP1-58188, Novus Biologicals); anti-Lamin B1 pAb (1:2,000, ab16048, abcam); anti-Ran mAb (1:2,000, 610340, BD Bioscience). To visualize the target molecules, cells were then incubated with Alexa Fluor 594-conjugated secondary antibody (1:2,000, Invitrogen, A32742). For amyloid fibril staining, cells were incubated with thioflavin S (ThS, 20 μg/mL in PBST) for 15 min and differentiated with 50% ethanol for 10 sec at room temperature. Counterstaining for nuclei was performed with DAPI (Thermo Fisher Scientific). Cells were then imaged and analyzed by the laser scanning confocal microscope as described above (see Live cell image analysis). For live cell imaging, tau reporter cells were cultured on μ-Dish 35 mm plate (81156, ibidi), seeded with pathogenic tau derived from TgTau^P301L^, and analyzed by live cell imaging. At 6 days post-seeding, time-lapse images of the cells were collected for 16-18 hours (10 min/frame for 96-108 frames) with Z-stack function under identical imaging settings. Image data were acquired with the laser scanning confocal microscope, ZEN Digital Imaging for LSM 700 (Zeiss) fitted with an environmental chamber at 37°C and 5% CO_2_ and analyzed using Zen 2010b SP1 imaging software (Zeiss) and Image J (https://imagej.nih.gov/ij/).

### Transmission electron microscopy (TEM)

Cells were collected and fixed in pre-warmed 2% paraformaldehyde in PB (0.1 M phosphate buffer, pH 7.3) for 20 min at 37°C and another 40 min at room temperature. The samples were post-fixed in 1% osmium tetroxide in PB for 1 hour and then incubated with 1% carbohydrazide in distilled water for 10 min at room temperature. After additional incubation with 1% osmium tetroxide for 1 hour, the samples were dehydrated in an ethanol series and infiltrated with an increasing concentration of Spurr’s resin (14300, Electron Microscopy Sciences) over several days. The infiltrated cell pellets were transferred to beam capsules and polymerized at 65°C for 24 hours. The resin-embedded pellets were sectioned with a thickness of 100 nm and incubated in 0.5% uranyl acetate for 1 hour at RT for negative staining. The thin sections on carbon grids were imaged using JEM-2100 LaB6 TEM (JEOL) with Gatan DigitalMicrograph (Gatan) software operated at 25 kV. TEM images were then analyzed using ImageJ software.

### Subcellular fractionation (Nuclear-cytoplasmic fractionation)

Cells were harvested after trypsinization and plated at 2×10^6^ cells/well of 6-well culture plates. On the next day, cells were cross-linked with 2% fresh formaldehyde (28908, Thermo Fisher Scientific) at 37°C for 10 min. The cross-linking reactions were quenched by adding the same volume of 1M glycine solution at 37°C for 5 min and cells were harvested. Nuclear and cytoplasmic extracts were prepared using NE-PER Nuclear and Cytoplasmic Extraction Reagents (78833, Thermo Fisher Scientific) following the manufacturer’s instructions. Briefly, cell membranes were disrupted by addition of the first detergent. Cytoplasmic extracts were recovered by centrifugation and the nuclei were then lysed with the second detergent to yield nuclear extracts. Nuclear Ran gradient was analyzed using capillary western assay. For reversal of the formaldehyde cross-links, the extracts were incubated with Fluorescent Master Mix (ProteinSimple) at 95°C for 20 min and analyzed by the capillary western assay. Extract purity was determined by probing with anti-β-tubulin pAb (NB600-936, Novus Biologicals), and anti-Lamin B1 pAb (ab16048, Abcam). For details, see western blot and capillary western assays below.

### Nuclear-cytoplasmic compartmentalization (NCC) assay

Cells were transfected with the NCC reporter construct which carries the IRES-linked sequences for GFP fused NES and RFP fused NLS under the control of EF1α promoter (pLVX-EF1alpha-2xGFP:NES-IRES-2xRFP:NLS) (Mertens et al., 2015). One μg of the construct was combined with 2 μL Lipofectamine 3000 (L3000-015, Thermo Fisher Scientific) and added to the cells. The cells were then incubated for 6 hours at 37°C and the media containing the DNA-liposome complex were replaced with fresh culture media. After 48 hours, images were obtained using the laser scanning confocal microscope as described above (see Live cell image analysis) and NCC were determined by Plot Profile analysis using Image J software.

### Fluorescence recovery after photobleaching (FRAP) analysis

For FRAP analysis of NCC, cells were plated on μ-Dish 35 mm plate and transiently transfected with NCC reporter construct (see NCC assay). On the next day, RFP signals in nuclear ROIs were obtained as time-lapse images (10 min/frame for 5 frames) (see Live cell image analysis) and then RFP were repeatedly bleached throughout the entire field. To determine recovery of RFP in nuclear ROIs, post-bleaching time-lapse images were collected for 6 hours (10 min/frame for 36 frames). Intensities of RFP in nuclear ROIs were measured using Image J software. For FRAP analysis of condensed liquid tau droplets, ES1 cells were plated on μ-Dish 35 mm plate and reference images were obtained. ROIs including NE tau inclusions were repeatedly bleached and time-lapse images were collected for 30 min (30 sec/frame for 55 frames).

### Sedimentation analysis

Sedimentation of tau in the seeded reporter cells was performed as previously described (Kaufman et al., 2016; Sanders et al., 2014) with some modifications. Briefly, clarified cell lysates were prepared as described above (see Limited proteolysis) and 10% of each lysate were set aside as total fractions. The rest were centrifuged at 100,000xg for 1 hour and the supernatants were placed aside as soluble fractions. The pellet was washed with 1.5 mL PBS prior to ultracentrifugation at 100,000xg for 30 minutes. For insoluble fractions, the pellet was re-suspended in RIPA buffer (50 mM Tris, 150 mM NaCl, pH 7.4, 1% NP-40, 0.5 % sodium deoxycholate, 4% SDS and 100 mM DTT) and sonicated at 30 amplitude for 3 min. Protein concentrations were normalized by BCA protein assay (Pierce) and tau in each fraction were analyzed by capillary western assay.

### Western blot and capillary western assays

Protein concentrations of each sample were normalized by BCA protein assay (Pierce). The samples were resolved on 15% Tris-Glycine gels or NuPAGE Bis-Tris gels (NP0343, Invitrogen), and transferred to PVDF membrane (Thermo Fisher Scientific). The membranes were blocked with 2% bovine serum albumin (BSA, Darmstadt) in TBST (TBS with 0.1% Tween 20) and probed with monoclonal (mAb) or polyclonal (pAb) antibodies at 4°C overnight: anti-tau mAb ET3 (Espinoza, de Silva, Dickson, & Davies, 2008) (1:500); anti-tau mAb RD4 (1:500, 05-804, Millipore); anti-Cleaved Caspase-3 pAb (1:2,000, #9661, Cell Signaling Technology); anti-Bax mAb (1:2,000, ab32503, abcam); anti-β-actin mAb (1:10,000, Abcam, ab20272). Anti-mouse IgG pAb conjugated to horseradish peroxidase (1:10,000, 170-6516, Bio-Rad) or anti-rabbit IgG pAb conjugated alkaline phosphatase (1:10,000, S3731, Promega) were used as secondary antibodies and visualized by detecting chemiluminescence (32209, Pierce) or fluorescence (S1000, Promega) signals. The membranes were stripped in western blot stripping buffer (46430, Thermo Fisher Scientific) and re-probed as needed.

Capillary western was performed as described in a previous report (Castle, Daude, Gilch, & Westaway, 2019). Reagents and equipment were purchased from ProteinSimple unless stated otherwise. Cell lysates or fractions were incubated with Fluorescent Master Mix at 95°C for 5 min. Four microliters of each sample were loaded into the top-row wells of plates preloaded with proprietary electrophoresis buffers designed to separate proteins of 12-230 kDa. Subsequent rows of the plate were filled with blocking buffer, primary and secondary antibody solutions, and chemiluminescence reagents, according to the manufacturer’s instructions. Primary antibodies were anti-tau mAb ET3 (Espinoza et al., 2008) (1:50), anti-Ran mAb (1:1,000, 610340, BD Bioscience), anti-β-tubulin pAb (1:1,000, NB600-936, Novus Biologicals), and anti-Lamin B1 pAb (1:1,000, ab16048, abcam). Secondary antibodies were anti-mouse or anti-rabbit secondary HRP conjugate. Peak area calculations and generation of artificial lane view were performed by the Compass software using the default Gaussian method.

### Limited proteolysis

Cell pellets were thawed on ice, lysed by triturating in PBS containing 0.05% Triton X-100 and protease inhibitors (cOmplete, Roche) and clarified by 5 min sequential centrifugations at 500xg and 1000xg. The cell lysates (1 μg/μL) were enzymatically digested with 50 μg/mL pronase E (Roche) at 37°C for 1 hour followed quenching with protease inhibitors and SDS-PAGE loading buffer, 15 μg/mL proteinase K (Ambion) at 37°C for 1 hour followed quenching with SDS-PAGE loading buffer, and 40 μg/mL thermolysin (Sigma) at 65°C for 30 min followed quenching with 0.5 M EDTA and SDS-PAGE loading buffer, respectively. The undigested tau fragments in each enzymatic reaction were determined by western blot analysis using anti-tau mAb ET3 (Espinoza et al., 2008) or anti-tau mAb RD4 (05-804, Millipore). For details, see western blot and capillary western assays above.

### Statistical analysis

The number of independent experiments or biological replicates of compared groups were at least n=3 for each observation. Statistical analysis for the quantitative data including cell viability, western blot, capillary western assay and FRAP analysis was performed using unpaired, two-tailed student t-test. Statistical analysis of all data was performed using PRISM version 5 software (GraphPad Software).

## Acknowledgements

Work in the Westaway lab was funded by the Canadian Institutes of Health Research (CIHR PS148962 and GER163048) and by Alberta Innovates Biosolutions (ABIBS AEP 201600021 and 20160023). Instrumentation was supported by the Canada Foundation for Innovation (NIF21633) and by the Alberta Synergies in Alzheimer’s and Related Disorders (SynAD) program, which is funded by the Alzheimer Society of Alberta and Northwest Territories through the ‘Hope for Tomorrow’ program and the University Hospital Foundation. DW was supported through a Canada Research Chair (Tier 1) and EG was supported by a scholarship from CONACYT (472481). We are indebted to the Neurological Tissue Bank of the Biobank-Hospital Clinic-IDIBAPS, Barcelona, Spain and Teresa Ximelis for sample and data procurement and to all brain donors and their families for generous brain donation for research. Special thanks go to Drs. Laura Molina-Porcel and Ellen Gelpi for the lamin B analyses presented in Table 1. The authors thank Dr. Xuejun Sun for assistance with EM image analysis and Dr. Valerie Sim for use of the LSM 710 microscope.

## Author contributions

S.G.K. and D.W. conceived the project. S.G.K., Z.Z.H., N.D., E.M., S.W., L.M.P. and E.G. performed experiments. All authors were involved in data collection and analysis. S.G.K. and D.W. wrote and revised the manuscript, which was approved by all authors before submission.

## Additional information

### Conflict of Interest

The authors declare that they have no conflict of interest.

## SUPPLEMENTARY MATERIALS

**Supplementary Figure 1.**
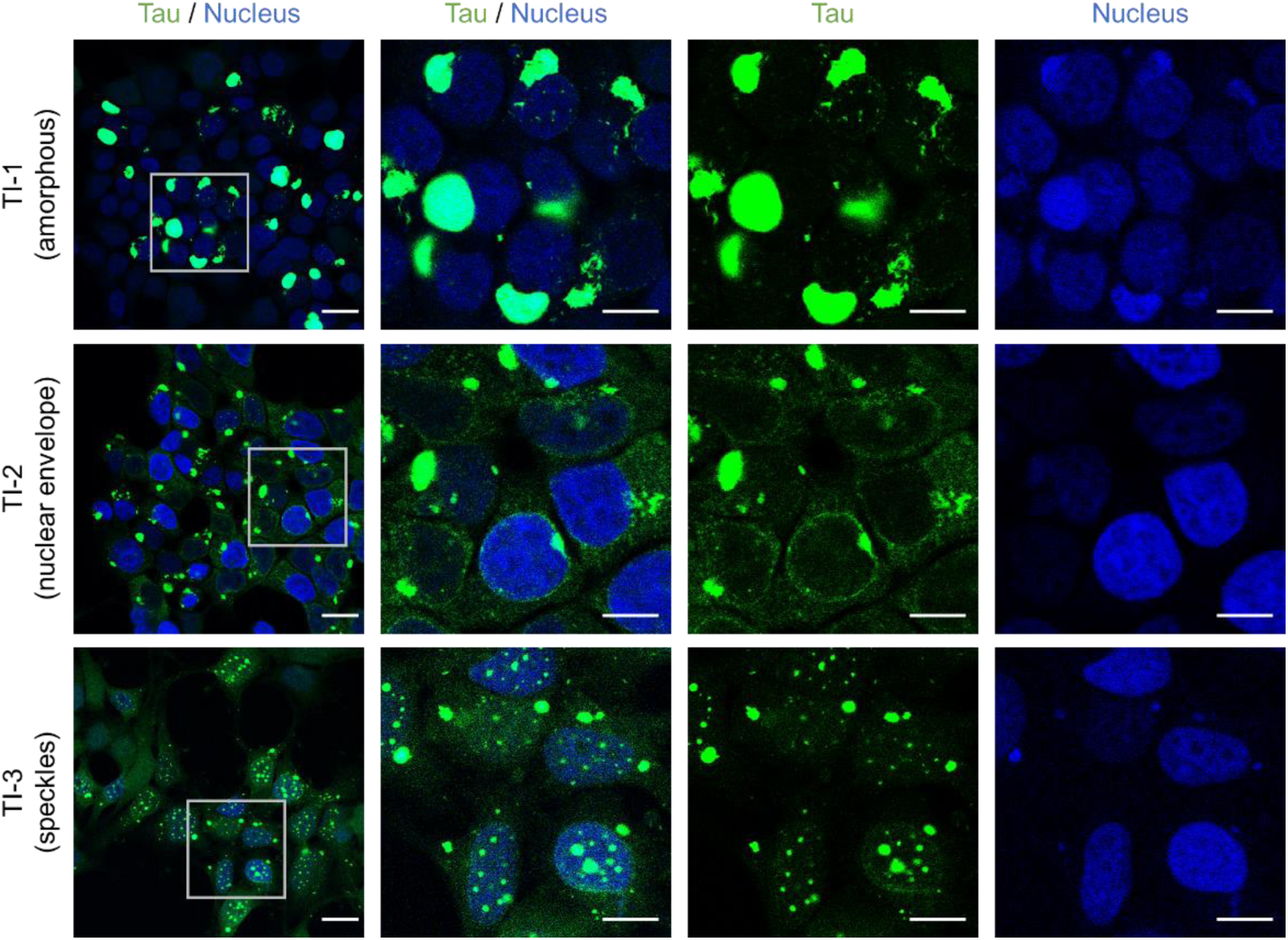
Tau seeding assay using tau reporter cells. The 4RD-YFP tau reporter cells (P301L/V377M) were seeded with tau derived from aged TgTau^P301L^ mice as per **Figure 2**. Tau inclusions were visualized using the YFP fusion tag (green) and nuclei were counterstained with DAPI (blue). Cytoplasmic and/or nuclear localization of various types of tau inclusion morphologies were verified with the nuclear staining. Scale bar, 20 μm and 10 μm in the boxed images. Related to **Figure 2**.

**Supplementary Figure 2.**
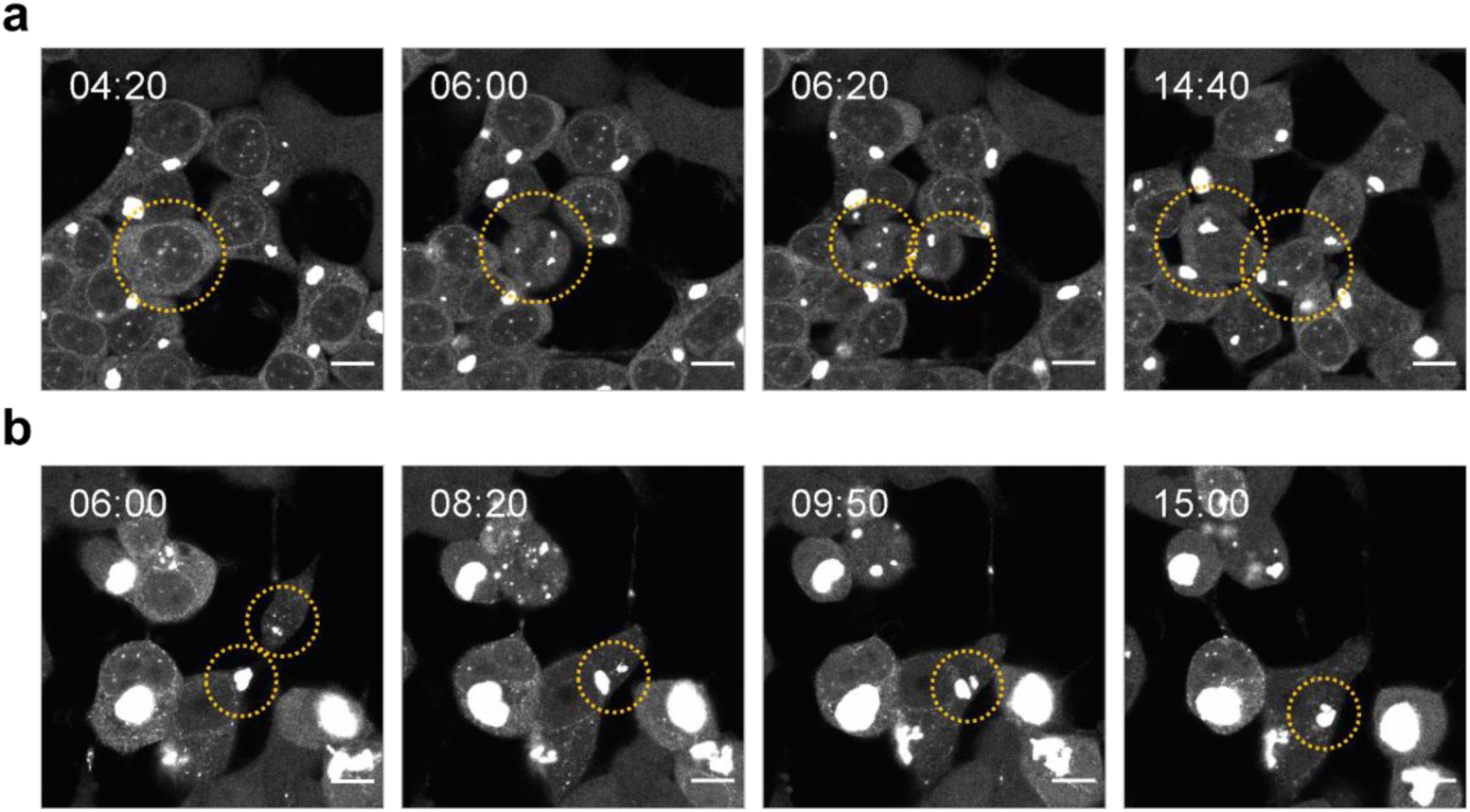
Live-cell imaging analysis of the seeded tau reporter cells. The 4RD-YFP tau reporter cells (P301L/V377M) were seeded with tau as per **Figure 2**. **a.** TI-positive cells underwent cell division and produced two daughter cells containing TIs (yellow cycles). **b.** TIs within cellular debris were absorbed by adjacent cells and combined with others, resulting in a bigger inclusion (yellow cycles). The cells were imaged every 10 min for 16 hours. Scale bar, 10 μm. Related to **Figure 2**.

**Supplementary Figure 3.**
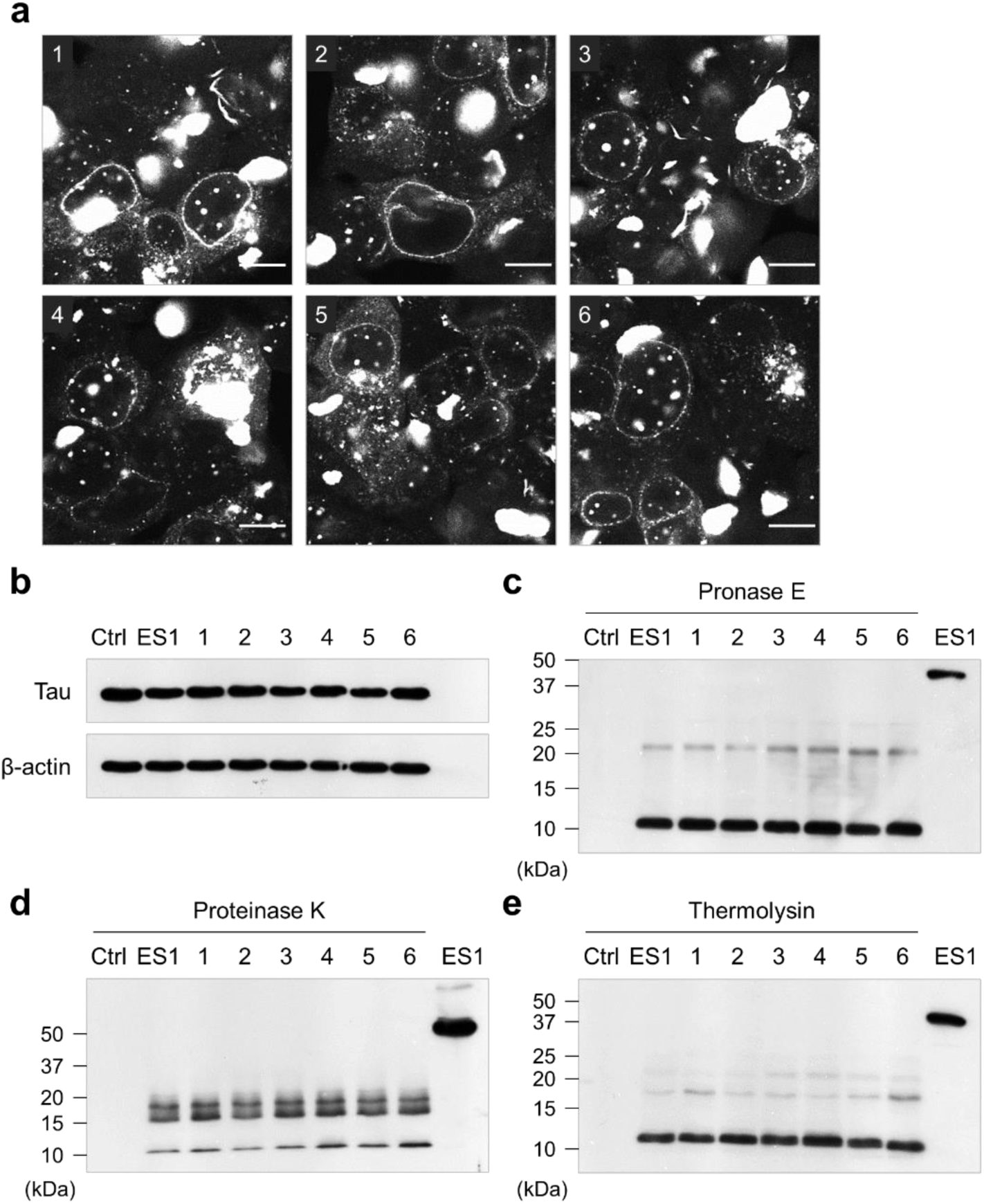
Heterogeneous tau inclusion morphology and limited proteolytic digestions. **a.** ES1 cells were re-subcloned by limiting dilution to obtain monoclonal cell populations representing each tau inclusion morphology. All six subclones (1 to 6) showed a heterogeneous phenotype the same as ES1 parental cells seen in **Figure 2a**. Scale bar, 10 μm. **b** to **e.** To differentiate the protected fibrillar cores of tau in individual cells, the cell lysates (**b**) were digested using pronase E (**c**), proteinase K (**d**), and thermolysin (**e**), and analyzed by western blot using anti-tau mAbs, ET3 and RD4. The limited proteolytic digestions revealed resistant core peptides in each subline (1 to 6) ranging from 10 to 25 kDa in size, while tau species in the reporter controls (Ctrl, 4RD-YFP P301L/V377M) were completely cleaved. The 10 kDa protease-resistant core appeared in all digestion conditions, and one or two bands between 15 to 20 kDa were shown depending on the enzymes tested. The patterns of the fragmented resistant cores were identical to each other, indicating that ES1 cells as well as the subclones reflect phenotypically similar monoclonal cell populations. Related to **Figure 3**.

**Supplementary Figure 4.**
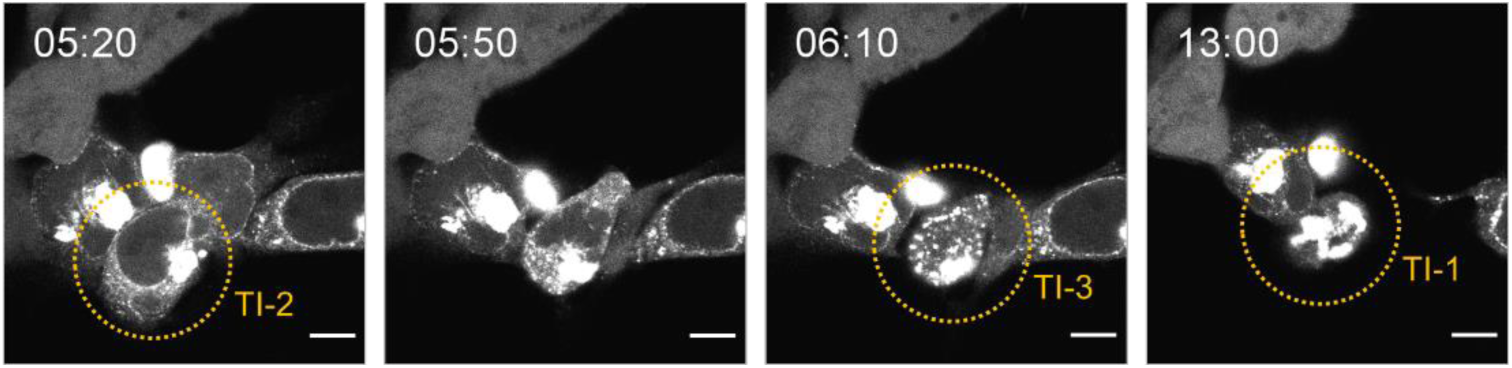
Morphological changes in tau inclusions. Live-cell imaging revealed that, in the seeded reporter cells (4RD-YFP P301L/V377M) as per **Figure 2**, TIs underwent morphological changes. TI-2 morphology (nuclear envelope, NE) was turned into TI-3 (speckles) and then TI-1 (amorphous). The cells were imaged every 10 min for 16 hours. Scale bar, 10 μm. Related to **Figure 3**.

**Supplementary Figure 5.**
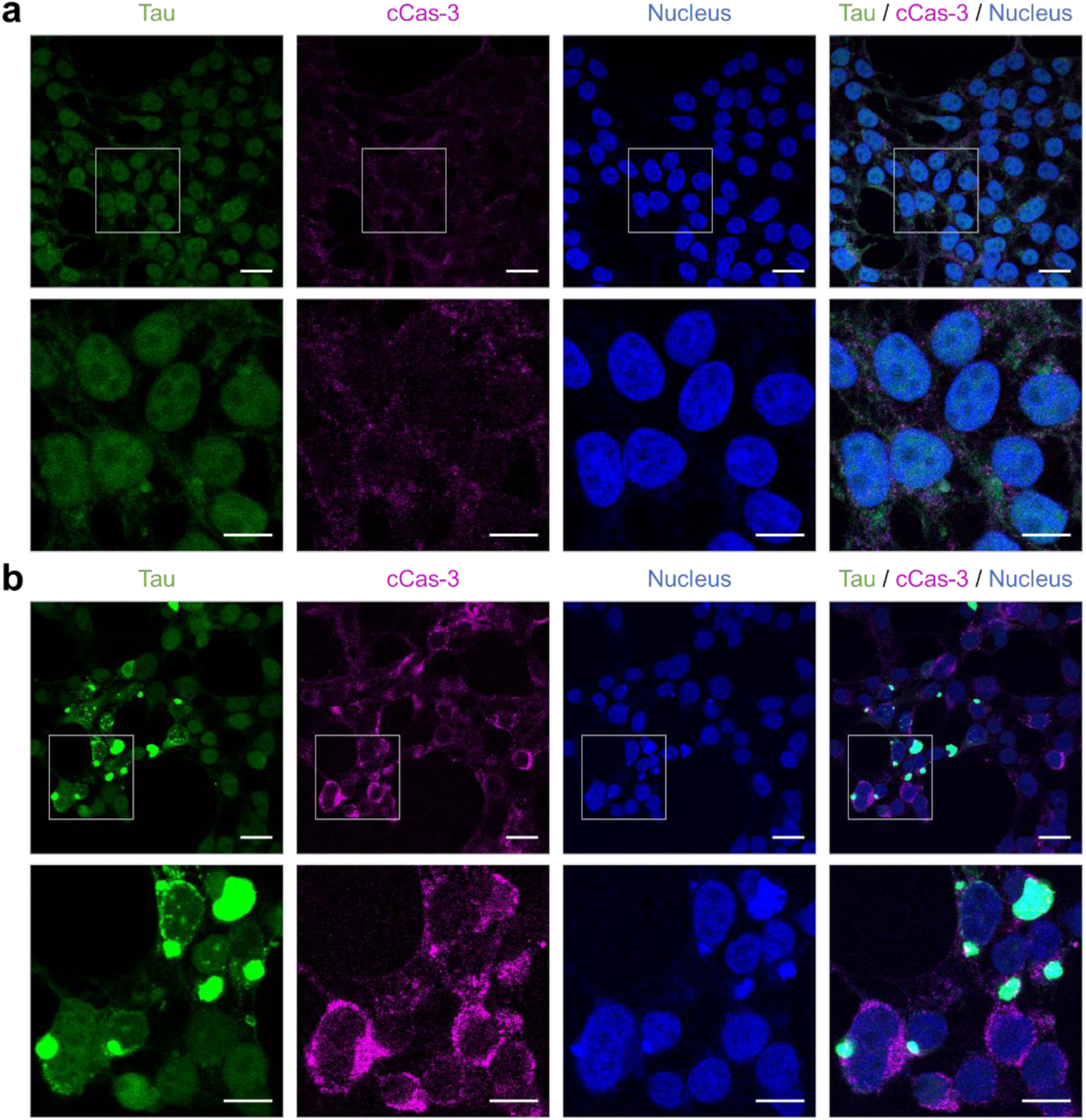
Increase in the level of cleaved Cas-3 with tau inclusions. Tau reporter cells (4RD-YFP P301L/V377M) were transiently seeded tau and performed immunocytochemistry for cCas-3 as per **Figure 2** and **Figure 5**. In comparison with control cells seeded with non-Tg brain homogenate (**a**), the level of cleaved Cas-3 was increased in the tau seeded cells and adjacent cells (**b**). Tau in green; cleaved Cas-3 in magenta; nuclei were counterstained with DAPI (blue). Scale bar, 20 μm and 10 μm in the boxed images. Related to **Figure 4**.

**Supplementary Figure 6.**
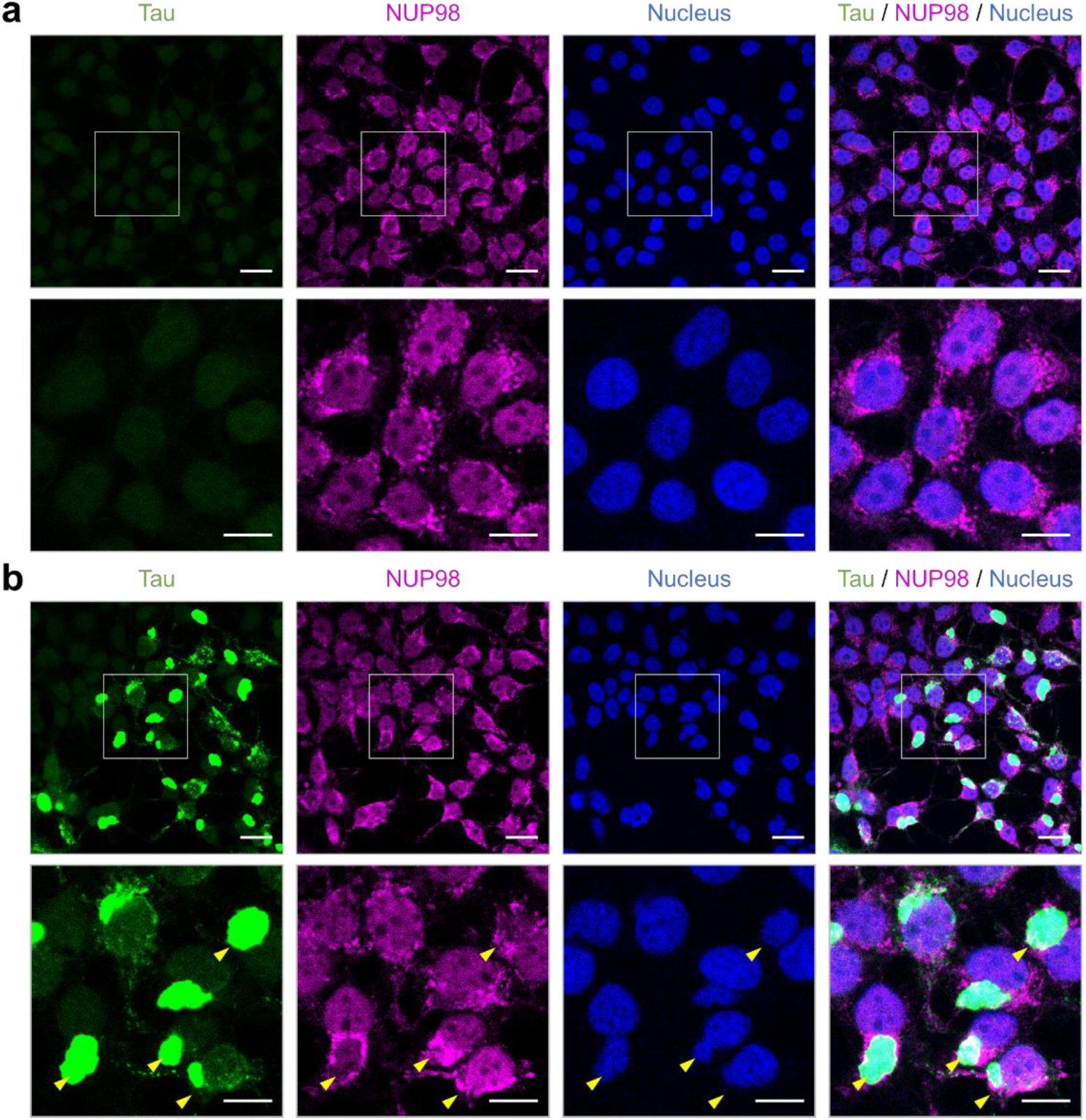
Mis-localization of NUP98 with tau inclusions. Tau reporter cells (4RD-YFP P301L/V377M) were seeded with tau and imaged as per **Figure 2** and **Figure 5**, respectively. NUP98 is one of the most abundant nucleoporins and contains Phe and Gly-rich repeats. In comparison with control cells seeded with non-Tg brain homogenate (**a**), mis-localization of NUP98 signals, which surrounded tau inclusions, were observed in the seeded cells (**b**). Tau in green; NUP98 in magenta; nuclei were counterstained with DAPI (blue). Scale bar, 20 μm and 10 μm in the boxed images. Related to **Figure 5**.

**Supplementary Figure 7.**
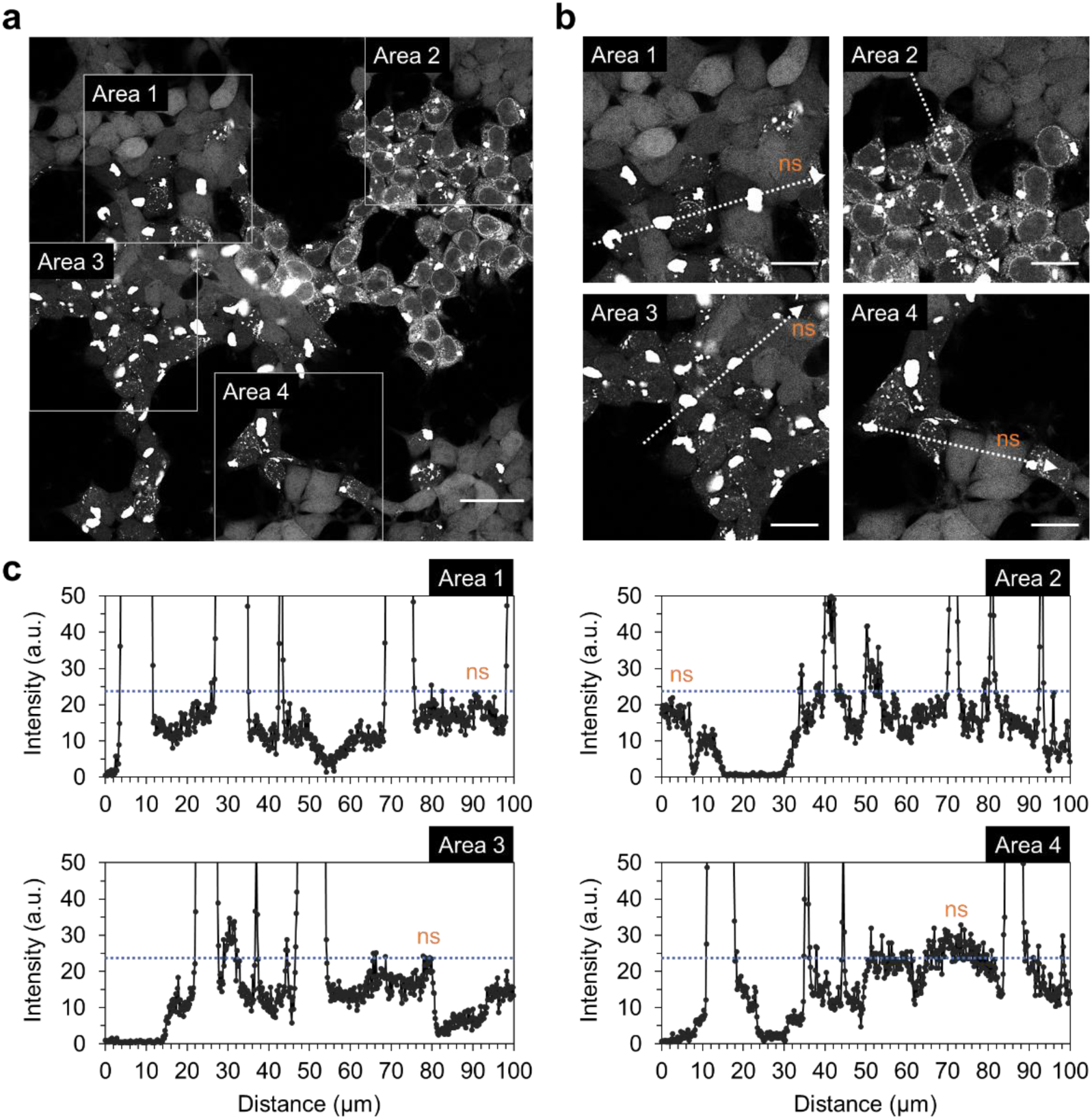
Condensation of tau-YFP into various inclusion morphologies. **a.** Transiently tau-seeded reporter cells (4RD-YFP P301L/V377M) as per **Figure 2** showed various inclusion morphologies. **b** and **c.** Plot profiling of the tau inclusions. Intensities of tau-YFP signals were measured from four different areas across the tau-positive and negative cells along the arrows with a length of 100 μm. a.u., arbitrary units; ns, non-seeded cells. Scale bar, 40 μm and 20 μm in the boxed images. Related to **Figure 8**.

## DESCRIPTION OF RELATED MANUSCRIPT FILES

File Name: **Supplementary Movie 1**

Description: Live cell imaging of tau reporter cells (4RD-YFP P301L/V377M) seeded with brain homogenate of clinically ill TgTau^P301L^ with CSA Type 2 profiling. Tau inclusions spread through cell division. Time-lapse movies were created at 6 days post seeding by recording photographs for 16 hours at one frame every 10 min (1/10 frame/min). Scale bar, 10 μm. Related to **Supplementary Figure 2a**.

File Name: **Supplementary Movie 2**

Description: Live cell imaging of the seeded tau reporter cells (4RD-YFP P301L/V377M) as per **Supplementary Movie 1**. Cell-to-cell spread of tau inclusions through membrane nanotubes. Time-lapse movies were created at 6 days post seeding by recording photographs for 16 hours at one frame every 10 min (1/10 frame/min). Scale bar, 10 μm. Related to **Figure 2c**.

File Name: **Supplementary Movie 3**

Description: Live cell imaging of the seeded tau reporter cells (4RD-YFP P301L/V377M) as per **Supplementary Movie 1**. Tau inclusions were adsorbed by adjacent cells. Time-lapse movies were created at 6 days post seeding by recording photographs for 16 hours at one frame every 10 min (1/10 frame/min). Scale bar, 10 μm. Related to **Supplementary Figure 2b**.

File Name: **Supplementary Movie 4**

Description: Live cell imaging of tau reporter cells (4RD-YFP P301L/V377M) as per **Supplementary Movie 1**. Multinucleated cells emerged by a failure in cell division. Time-lapse movies were created at 6 days post seeding by recording photographs for 16 hours at one frame every 10 min (1/10 frame/min). Scale bar, 10 μm. Related to **Figure 2d**.

File Name: **Supplementary Movie 5**

Description: Live cell imaging of tau reporter cells (4RD-YFP P301L/V377M) as per **Supplementary Movie 1**. NE tau inclusions were transformed into speckle and then amorphous shapes. Time-lapse movies were created at 6 days post seeding by recording photographs for 16 hours at one frame every 10 min (1/10 frame/min). Scale bar, 10 μm. Related to **Supplementary Figure 4**.

File Name: **Supplementary Movie 6**

Description: Live cell imaging of tau reporter cells (Dox:GFP-0N4R P301L) as per **Supplementary Movie 1**. NE tau inclusions were transformed into amorphous shapes. Time-lapse movies were created at 6 days post seeding by recording photographs for 16 hours at one frame every 10 min (1/10 frame/min). Scale bar, 10 μm. Related to **Figure 3b**.

File Name: **Supplementary Movie 7**

Description: Live cell imaging of tau reporter cells (4RD-YFP P301L/V377M) as per **Supplementary Movie 1**. Multinucleated reporter cells containing NE tau inclusions underwent apoptotic cell death. Time-lapse movies were created at 6 days post seeding by recording photographs for 16 hours at one frame every 10 min (1/10 frame/min). Scale bar, 10 μm. Related to **Figure 4g**.

File Name: **Supplementary Movie 8**

Description: Live cell imaging of tau reporter cells (4RD-YFP P301L/V377M) transiently transfected with NCC reporter construct. For FRAP analysis, 5 reference photographs were taken at the beginning and RFP were photobleached. Time-lapse movies were created by recording photographs for 6 hours at one frame every 10 min (1/10 frame/min). Scale bar, 10 μm. Related to **Figure 7b**.

File Name: **Supplementary Movie 9**

Description: Live cell imaging of ES1 cells transiently transfected with NCC reporter construct as per **Supplementary Movie 8**. Time-lapse movies were created by recording photographs for 6 hours at one frame every 10 min (1/10 frame/min). Scale bar, 10 μm. Related to **Figure 7b**.

File Name: **Supplementary Movie 10**

Description: Live cell imaging of NE tau inclusions in ES1 cells. For FRAP analysis, 5 reference photographs were taken at the beginning and NE tau inclusions were photobleached. Time-lapse movies were created by recording photographs for 30 min at one frame every 30 sec (1/30 frame/sec). Scale bar, 10 μm. Related to **Figure 8c** and **8d**.

File Name: **Supplementary Movie 11**

Description: Live cell imaging of multinucleated cells containing NE tau inclusions (ES1 cells). For FRAP analysis, 1 reference photograph was taken at the beginning and NE tau inclusions were photobleached. Time-lapse movies were created by recording photographs for 30 min at one frame every 30 sec (1/30 frame/sec). Scale bar, 10 μm. Related to **Figure 8e**.

## Notes

### Competing Interest Statement

The authors have declared no competing interest.

